# OsIDS3L, a Non-Canonical Dioxygenase Enhancing Iron Homeostasis and Micronutrient Biofortification in Rice

**DOI:** 10.64898/2025.12.07.692860

**Authors:** Jyoti Aggarwal, Kuo-Chen Yeh

**Author notes:** corresponding author Kuo-Chen Yeh.

## Abstract

- Graminaceous plants synthesize some organic compounds known as phytosiderophores (PSs) for iron (Fe) uptake. The first PS, deoxymugineic acid (DMA), is common to all grasses. However, PSs synthesized from DMA, hydroxylated PSs, have higher efficiency for Fe chelation and are species-specific. Hydroxylated PS, mugineic acid (MA) synthesis via the Fe(II)-2-oxoglutarate-dependent dioxygenase enzymes HvIDS2 and HvIDS3 have been reported in barley.
- Here, we report that IDS3-Like (OsIDS3L), a homolog of the MA biosynthesis gene, is responsive to Fe deficiency in rice. Functional analysis using overexpression (OE) lines demonstrates OsIDS3L’s role in Fe-deficiency tolerance and micronutrient assimilation in seeds. However, MA was not detected, indicating functional divergence of IDS in rice. Increased DMA and NA levels in OE lines support enhanced Fe uptake and homeostasis.
- CRISPR-generated mutants lack an obvious phenotype and show no substrate DMA accumulation. Expression analysis suggests regulation of Fe-responsive genes by both gain- and loss-of-function lines. A computational metabolomics analysis, supported by chemotype justification, suggests a role for OsIDS3l in an enzyme reaction upstream of DMA.
- The results not only identified a gene potential for agronomy-specific traits and biofortification, but also shed light on a possible non-canonical PS biosynthesis pathway.

## Introduction

Iron (Fe) is an essential micronutrient for plant growth and development (Briat, Dubos and Gaymard, 2015). However, it is present in the insoluble ferric [Fe(III)] form in aerobic, alkaline soil, which is poorly bioavailable to plants (Lindsay and Schwab, 1982; Slessarev *et al*., 2016). Graminaceous plants have evolved a unique strategy for Fe uptake and overcome this stress, in which plant roots synthesize and secrete low-molecular-weight organic compounds in the rhizosphere known as Phytosiderophores (PSs). These PSs chelate Fe(III) in the rhizosphere, forming PS-Fe(III) complexes that are readily taken up by the plant root transporters. To date, nine PSs have been identified, with molecular weights ranging from 294 to 336 Da (Takagi, 1976; Hiradate, Ma and Matsumoto, 2007). These PSs possess six coordination sites-three carboxyl, two amine, and one hydroxyl group-that facilitate Fe(III) chelation (TAKEMOTO *et al*., 1978; Sugiura *et al*., 1981).

PSs are synthesized via a series of enzymatic reactions originating from methionine. One key intermediate in this pathway is nicotianamine (NA), which is ubiquitous in plants and plays a central role in intracellular metal homeostasis (Hiradate, Ma and Matsumoto, 2007). NA is further converted into deoxymugineic acid (DMA), the primary PS produced by all graminaceous plants (Bashir *et al*., 2017). Some graminaceous plants like barley, rye and oats catalyze hydroxylation of DMA at specific C-positions, yielding structurally diverse PSs, known as hydroxylated PSs. These hydroxylated PSs are correlated with enhanced Fe-chelating efficiency and confer Fe-deficiency tolerance in plants (Römheld and Marschner, 1990; Von Wiren, Khodr and Hider, 2000). However, plants like rice and maize, exhibit greater sensitivity due to PS production in both qualitative and quantitative terms (Römheld and Marschner, 1990).

Rice sensitivity to Fe deficiency, particularly under aerobic and alkaline soil conditions, is a major constraint on yield (Lindsay and Schwab, 1982; Fan *et al*., 2012; Slessarev *et al*., 2016; Carrijo, Lundy and Linquist, 2017; Calabrese and Porporato, 2019; Kanwar, Baby and Bauer, 2021; Bakaram *et al*., 2022). This soil condition-derived sensitivity is reported to be mitigated by (1) exogenous Fe application (Pal *et al*., 2008; Kumar *et al*., 2019; de Campos Carmona *et al*., 2021; Sreelakshmi *et al*., 2021; Xu *et al*., 2021; Kumar and Yer, 2023), (2) transgenic approaches that enhance PS biosynthesis (Connorton and Balk, 2019; Masuda *et al*., 2019). This highlights the significance of Fe acquisition capacity for sustainable cultivation in challenging conditions.

The biosynthesis of MA in barley is mediated by Fe(II)-2-oxoglutarate–dependent dioxygenases (2OGDs), HvIDS2 and HvIDS3 (Iron Deficiency Specific). HvIDS2/3 were originally identified through differential hybridization experiments under Fe-deficient conditions and IDS1 encoded a metallothionine (Nakanishi *et al*., 2000). Homologous sequence of HvIDS2/3 has since been identified only in *Secale cereale* and wheat (Nakanishi *et al*., 2000; Mathpal *et al*., 2018). This raises the question of whether rice possesses the capacity to synthesize hydroxylated PSs. With the availability of sequenced genomes and improved databases, we revisited this question and identified a rice gene Os07g07410 with notable sequence similarity to HvIDS3. We refer to the gene as IDS3-like (OsIDS3L) and hypothesize that it may encode a functional enzyme in rice’s Fe-deficiency response.

In this study, we functionally characterized OsIDS3L through a combination of functional studies and metabolic profiling. We reported OsIDS3L as a single-copy gene in rice that enhances Fe uptake and homeostasis in overexpression lines, conferring beneficial agronomic and biofortification traits. We also studied the CRISPR-generated mutant phenotype. Our results reveal that OsIDS3L plays a distinct role from HvIDS3. Despite synthesizing MA, it modulates the accumulation of PS biosynthesis compounds and enhances rice Fe acquisition. Using computational metabolomics, we speculate OsIDS3L enzyme activity in a possible non-canonical pathway of DMA synthesis. We have discussed OsIDS3L’s role in developing Fe-efficient and nutritionally improved rice varieties, and proposed a model for its enzymatic activity.

## Methods and Methodologies

### Homology analysis/Bioinformatics analysis

Homologous gene was identified using BLAST search at different databases like NCBI, Ensemblplants, Phytozome, RAP-DB, MSU-RGAP, and Kitbase (Altschul *et al*., 1990; Goodstein *et al*., 2012; Kawahara *et al*., 2013; Sakai *et al*., 2013; Kersey *et al*., 2014; Li *et al*., 2017; Hamilton, Li and Buell, 2025). The multiple sequence alignment and pairwise sequence similarity were performed using QIAGEN CLC Genomics Workbench 11. Protein structure of HvIDS3 (Q40062) and OsIDS3L (Q69LD9) was prediction using Alphafold (Jumper *et al*., 2021). The superimposition of protein structure was performed using PyMol (Schrödinger, no date) and the TM-alignment score was calculated using TM-align (Zhang and Skolnick, 2005).

### Phylogenetic analysis

Putative 2OGD for rice were retrieved from earlier study and the methodology was used to retrieve and classify 2OGDs for barley (Table S1a-b) (Kawai, Ono and Mizutani, 2014). Briefly, the 2OGD proteins for barley were detected from whole-protein sequences. The whole amino acid sequences of barley (*Hordeum vulgare* Morex V3) were retrieved from the Phytozome database (Goodstein *et al*., 2012). Since 2OGD proteins with 2OG-FeII_Oxy motif (Pfam id, PF03171 (Aravind and Koonin, 2001)) is well conserved and characterized, we regarded proteins containing one 2OG-FeII_Oxy motif as a putative 2OGD. Therefore, we performed a motif search analysis using HMMER at CLC Genomics Workbench 11, with a cut-off e–value of 2e−4. Since we implied motif search over pairwise similarity search with BLAST to identify 2OGD in barley, the putative 2OGDs may include distantly related proteins. Therefore, we performed amino acid sequence alignment using MUSCLE over CLUSTAL OMEGA (Edgar, 2004). Further, JTT model was used for amino acid substitution and maximum-likelihood phylogenetic tree was generated using MEGA11 software. The trees constructed in this study are unrooted.

### Plant materials and growth conditions

*Oryza sativa* Kitaake was used as wild type and the background for genetic transformations in this study. *Hordeum vulgare* seeds were used to amplify barley CDS. Rice seeds were surface sterilized using 50% Sodium hypochlorite for 30 minutes and were washed with sterile distilled water 4-5 times for 5 minutes. Seeds were then germinated on half strength Murashige and Skoog culture medium in presence or absence of NaFeEDTA for control and Fe deficient media respectively at 28°C in day and 25°C at night under a long-day photoperiod (16h day/8h night). MES was used as a buffering agent with a concentration of 0.5g/l and the pH were maintained at 5.5. To generate alkaline condition, HEPES was used as buffering agent and the media pH was modulated to 10 using NaOH. An agar concentration of 8g/l was used for solidifying media unless mentioned. High agar percentage was implied to test the drought conditions (Gonzalez *et al*., 2023, 2024). Same media was used as hydroponics media without agar for longer phenotypic observation and for ICP-OES sampling.

For soil experiments, 10-14d old seedlings were transferred to greenhouse with a long photoperiod (16h day/8h night) and temperature maintained at 28/ 25°C in day/ night respectively. The water-logging and water-saving conditions were modulated by maintaining water level in the tank where pots are kept. Water level until the rice shoot submerged to 4-5 cm (pots submerged in water) was considered water-logging condition whereas when water is kept for 5-10 cm in tank containing pots was implied as water-saving condition. Pots used in the study have a hole in the middle. The weeds were eradicated weekly to remove bias of nutrient distribution among the rice and weeds. The experiment was repeated twice in greenhouse and once in the rice field. For the field experiment, alternate wet and dry method of watering was implied.

Rice seedlings at different stages were harvested for the respective experiments of expression analysis, ICP-OES and metabolomics analysis. To measure the physiological parameters like chlorophyll content, chlorophyll fluorometer (OptiSciences, CCM-300 Chlorophyll Content Meter) was used.

### Elemental analysis

Elemental analysis was performed using Inductively Coupled Plasma-Optical Emission Spectrometry (ICP-OES). Samples are washed with 10mM calcium chloride for 20 minutes and consecutive washing with double distilled water. Thereafter samples were dried at 70 °C in oven for 2-3 days until complete dry. 20-100mg of sample is weighed for one sample in Teflon tubes, and 2:1 (1ml: 0.5ml) ratio of nitric acid and hydrogen peroxide was added for microwave digestion at 160 °C using MARS 6TM PFAS Extraction System (CEM Corporation, Matthews, NC, USA). The digested sample was then transferred to 15ml falcon tube and 8.5 ml of 2% nitric acid was added. The solution was filtered using 0.45µm filters to new falcon and stored until elemental analysis.

### Construction of transgenic materials

To generate over-expression lines, the full-length coding sequence of *OsIDS3L* and *HvIDS3* was amplified from the cDNA of root samples of *Oryza sativa L. japonica*. cv. Kitaake and *Hordeum vulgare.* The cDNA was then cloned using TA cloning vector; pCR8/GW/TOPO, sequenced and confirmed. The entry clone generated was then used for LR gateway recombination reaction in P_UBQ_C-GFP comprising C-terminal GFP (Grefen *et al*., 2010).

To generate the reporter lines, we first studied the gene structure. OsIDS3L (Os07g07410) is at the positive strand of chromosome 7 and 562 bp downstream is Os07g07400 at the negative strand. Therefore, 562 bp promoter region was amplified and similarly cloned to entry vector; pCR8/GW/TOPO which further facilitated LR gateway cloning to pGWB3 vector comprising GUS sequence (Nakagawa *et al*., 2007).

To generate CRISPR mutants, sgRNA are designed using the http://crispr.hzau.edu.cn/cgi-bin/CRISPR2/CRISPR online tool, and selected based upon the secondary structure of sgRNA, maximum on-target efficiency, and minimum off-target efficiency. The pYLCRISPR/Cas9PUBQ-H-based constructs are created using Gibson assembly as previously reported (Ma *et al*., 2015). Four guide RNAs (gRNAs) targeting different exons of *OsIDS3L* were employed to enhance editing efficiency. Transgenic plants were screened to identify Cas9-free homozygous mutants, and three independent KO lines carrying distinct mutations were selected for experiments. Similarly, dmas1 mutant was also generated in Kitaake background as a positive control by targeting Os03g13390.

All the generated destination vectors were transformed into *Agrobacterium* strain EHA105 for the transformation of rice immature embryogenic callus culture generated on N6D media using the *Agrobacterium*-mediated callus transformation method as discussed earlier (Toki *et al*., 2006). The successfully transformed callus were screened on appropriate antibiotic media and regenerated. The seeds of the transgenic Kitaake plants were acquired from Tainan greenhouse. The overexpression lines, and the reporter lines were confirmed of the transgene and were screened and propagated for homozygous T3 lines. The mutants were screened by sequencing the Polymerase Chain Reaction (PCR) products spanning gRNA. Mutations were detected by aligning the sequence chromatograms of these PCR products with those of the WT controls. Several individual homozygous or heterozygous plants were obtained, and three homozygous lines were selected for further experiments.

### Subcellular localization

To study the subcellular localization, the P_UBQ_C-GFP constructs generated for transgenic construction were implied. Rice protoplasts were subsequently prepared from 7 day old wild-type rice shoot issues. Protoplasts were then transformed with empty vector (EV-P_UBQ_C:GFP control) as a negative control, P_UBQ_:*OsIDS3L*:*GFP* and P_UBQ_:*HvIDS3L*:*GFP* (as a positive control) constructs. The transformed protoplasts were observed under the confocal microscope after 12-14 hours of transformation at 488nm/ 505-530nm for GFP excitation and emission spectra.

### Histochemical staining

The generated transcriptional fusion lines were implied for GUS staining as described earlier (Bashir *et al*., 2006). Briefly, T1 transgenic lines were grown on ½ MS for different days under iron-sufficient (+Fe; 50 µM NaFeEDTA) and iron-deficient (–Fe; no added Fe) conditions. Whole seedlings were incubated in GUS reaction buffer (1 mM X-Gluc, 50 mM sodium phosphate pH 7.0, 0.5 mM K₃[Fe(CN)₆], 3 mM K₄[Fe(CN)₆], 20% methanol) at 37 °C after gentle vacuum to the plants for 30 minutes.

### RNA isolation and quantification of gene expression

Samples were harvested and snap chilled immediately. Total RNA was extracted from the rice samples using Trizol reagent according to the manufacturer’s instructions. Genomic DNA (gDNA) was removed and the first-strand complementary DNA (cDNA) was synthesized via Qiagen cDNA synthesis kit. RT-PCR was performed to amplify full length coding sequence (CDS). SYBR green was used for RT-qPCR analysis.

### RNA-seq analysis

Plants were germinated on half MS with and without 50µM NaFeEDTA for 8 days. Root samples were harvested and frozen in liquid nitrogen. Samples were stored at -80° C until RNA extraction. Total RNA was extracted using *mir*Vana miRNA Isolation Kit (Thermo Fisher Scientific Inc., Wilmington, DE, USA). RNA quality and concentration was analyzed using NanoDrop 1000 UV-Vis Spectrophotometer (Thermo Fisher Scientific Inc., Wilmington, DE, USA). Sequencing libraries were generated and sequenced on NovaSeq 6000 platform (Illumina Inc., CA, USA). Raw sequence libraries were quality filtered to remove adapter and low-quality bases using Phred scores. High quality reads were trimmed and aligned to reference genome of Oryza sativa using Hisat2. The best match was used to summarize mapped reads to gene count data. Expression values were derived from FPKM values (fragments per kilobase of exon model per million mapped reads). DESeq2 normalized values are used to report the expression differences.

### Phytosiderophore extraction from the root samples

Metabolic experiments are performed mostly on young seedling’s roots which helps to reduce bacterial influence to plants. Further, the bacterial metabolites would make unsure of the compound origin. Therefore, all Phytosiderophores (PSs) related experiments were performed in sterile growth conditions in agar media. Plants were germinated and grown on respective conditions and PSs were extracted from the snap-chilled root samples. The sample was grinded using mortar pestle in liquid nitrogen, and was dried in vacuum freeze-dryer (FD24-20PL, KingMech, New Taipei, Taiwan; at −60 to −80°C for 8-10h). The 50mg sample was weighed and 1ml of 80% methanol was added, vortexed and sonicated for 30 min at 4°C to extract the PSs. The mixture was centrifuged and clear solution was used as extract for LC-ESI-Q-TOF-MS/MS analysis. 2 ppm fenclonine (4-Chloro-DL-phenylalanine, Sigma C6506, CAS: 7424-00-2) was added as internal standard while extracting the metabolites.

### LC-ESI-Q-TOF-MS/MS experiment

The samples were separated using an Agilent 1290 II Infinity Ultra-High-Performance Liquid Chromatography system (Agilent Technologies, Palo Alto, CA, USA) and detected using Agilent 6545XT quadrupole time-of-flight (Q-TOF) mass spectrometer (Agilent Technologies, Palo Alto, CA, USA) equipped with an Agilent Jet-stream source.

The samples were separated by using ACQUITY UPLC T3 column (1.7 μm, 2.1 × 100 mm, Waters Corp., Milford, MA, USA). The column temperature was maintained at 30°C. The mobile phases were double-distilled water (eluent A) with 0.1% formic acid and 70% acetonitrile: water with 20 mM ammonium acetate (pH 3.5) (eluent B). The flow rate was 0.2 mL/min and the injection volume were 5 uL.

The instrument was operated in positive full-scan mode and collected from a m/z of 100–1100. The MS operating conditions were optimized as follows: Vcap voltage, 4.0 kV; nozzle voltage, 0.25 kV; nebulizer, 45 psi; gas temperature, 200° C; sheath gas temperature, 300° C; sheath gas flow (nitrogen), 10 L/min; drying gas flow (nitrogen), 8 L/min. The MS/MS data acquisition mode was set to auto MS/MS. Fixed collision energies of 20.0 was chosen at a scan rate of 4.0 spectra/s. The chromatogram acquisition, detection of mass spectral peaks, and their waveform processing were performed using Agilent LC/MS Data Acquisition software 9.0, Agilent Qualitative Analysis 10.0 and Agilent Profinder 10.0 software (Agilent, USA).

### Untargeted metabolomics analysis

Untargeted metabolomics was performed for the three genotypes (WT, OE and ko) germinated and grown under control (C, Fe-sufficient) and treated (T, 0Fe) conditions for 8 days. The Agilent resulting file (in .d format) was converted to mzML format using MSConvert tool within ProteoWizard Library (*ProteoWizard: Home*, no date; Kessner *et al*., 2008; Martens *et al*., 2010). The features were then curated using MZmine (Schmid *et al*., 2023) software and blank subtracted on FBMNstats (Shah *et al*., 2023). The MZmine curated features and spectra file was used for feature based molecular networking (FBMN) at Global Natural Product Social Molecular Networking (GNPS) with cosine score of 0.7 (Nothias *et al*., 2020). The unannotated features were further annotated using SIRIUS, CSI:FingerID and Zodiac applications (Dührkop *et al*., 2019). The visualization of network was facilitated by Cytoscape (Shannon *et al*., 2003) and were analyzed for edges with a most probable m/z difference (15.995; for catalyzed by a Fe(II)-2OG dioxygenase. The statistical analysis was performed on MetaboAnalyst 6.0 (Pang *et al*., 2024). Data were log2 transformed to stabilize variance across the dynamic feature ranges. Data were mean centered and auto-scaling was performed to reduce variance and the emphasis on very low variance features. Metabolic modification was determinded using Modifinder (Shahneh *et al*., 2024).

## Results

### Os07g07410, OsIDS3L is a single-copy rice gene homologous to barley IDS2/IDS3

Comparative genomics using BLAST identified Os07g07410 as a likely ortholog of the barley iron deficiency specific dioxygenase, HvIDS2 and HvIDS3. A dominant transcript Os07g07410.2 encodes a protein of 340 amino acids that shares 60% identity with HvIDS3 and 48% identity with HvIDS2 (Fig. 1a-c, Fig. S1). Multiple sequence alignment demonstrates conserved domains including DIOX-N (amino acids 41–152; Pfam ID: PF14226) and the Fe(II)-2OG dioxygenase domain (amino acids 192–291; Pfam ID: PF03171) along with conserved motifs, Fe(II) binding HX(D/E)XnH motif and 2OG-binding YXnRXS motif (Kawai, Ono and Mizutani, 2014) (Fig. 1a-c). Phylogenetic analysis of Fe(II)-2OG dioxygenases from rice and barley placed Os07g07410 in the same clade as HvIDS2/3, excluding other rice 2OGDs (Fig. S2). The monophyletic ‘IDS clade’ likely represents lineage-specific homologs involved in PS metabolism. Notably, rice has only one gene Os07g07410 in the IDS clade, whereas barley has multiple paralogs as reported earlier (Aleksza *et al*., 2024). An EnsemblPlants gene family analysis further confirmed that Os07g07410 is a single-copy gene (Fig. S3). Based on these results, we designated *Os07g07410.2* as OsIDS3L, the *HvIDS3*-Like gene in rice.

**Fig. 1.**
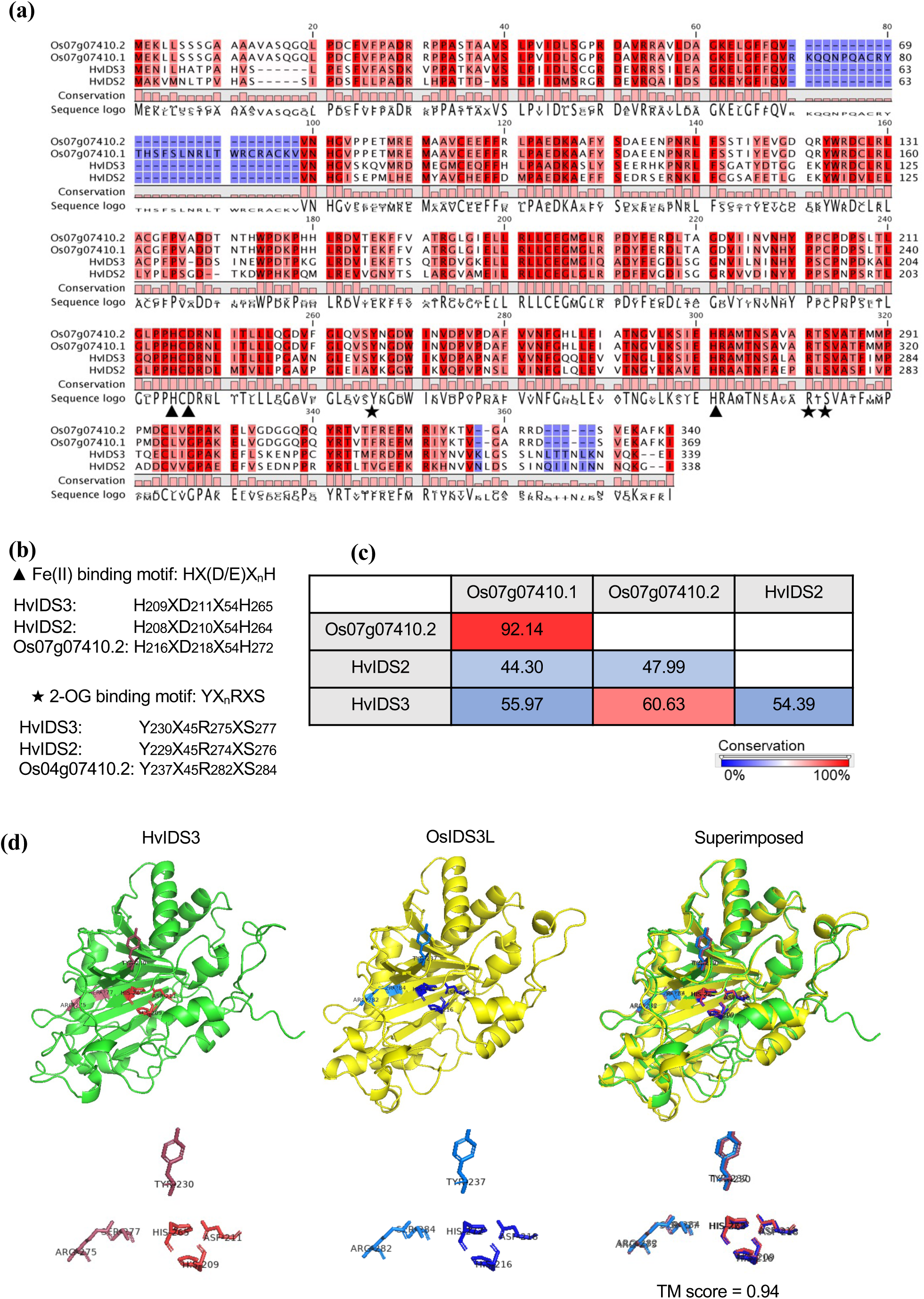
Os07g07410 is a single copy gene with shared homology to HvIDS2 and HvIDS3. **(a)** Multiple protein sequence alignment of HvIDS2, HvIDS3 and Os07g07410. Alignment conservation is color coded from blue (0%) to red (100%). Conserved Fe(II)-binding HX(D/E)XnH triad motifs and 2-OG binding YXnRXS motifs are marked by black triangles and stars respectively. **(b)** Expanded view showing the amino acid positions of the conserved motifs in the aligned sequences. **(c)** Pairwise protein sequence similarity (in percentage) among HvIDS2, HvIDS3, and Os07g07410. **(d)** Superimposed alfafold predicted 3D structure of HvIDS3 (green) and OsIDS3L (yellow) aligned and visualized in PyMOL. The conserved Fe(II) binding motif and 2-OG binding motifs are highlighted in shades of red and blue. The superimposed structures (α-helices and β-strands with conserved motifs) support structural homology of OsIDS3L with HvIDS3.

The function of an enzyme corresponds to the protein structure (Ribeiro *et al*., 2023). Therefore, we performed structural modelling using PyMOL. The Alphafold-predicted 3D structure of OsIDS3L aligned closely with HvIDS3, with a high TM-score of 0.95, indicating a highly similar spatial configuration and, likely, comparable catalytic function (Zhang and Skolnick, 2005; Jumper *et al*., 2021). Nonetheless, some structural differences were observed. OsIDS3L contains 15 α-helices and 11 β-strands, whereas HvIDS3 contains 11 α-helices and 11 β-strands (Fig. 1d). This altogether suggests that Os07g07410.2 is a functional homolog of HvIDS3, hereafter OsIDS3L, and may be involved in phytosiderophore biosynthesis and iron homeostasis.

### *OsIDS3L* is expressed in roots and reproductive tissues and is induced by Fe deficiency

To examine the expression pattern of OsIDS3L in rice, we mined public datasets showing OsIDS3L expression in roots and panicles (Fig S4). To extend the observation, we generated OsIDS3L promoter-driven GUS, P_OsIDS3L_: GUS, transgenic lines. Histochemical GUS staining revealed strong promoter activity in roots of seedlings. OsIDS3L expression was observed in both Fe-sufficient and Fe-deficient conditions. Besides, the GUS expression was detected in anthers of flowers at different stages (Fig. 2a). This aligns with previous reports where MA is synthesized from barley’s anther-derived suspension cultures (Nishizawa *et al*., 1989).

**Fig. 2.**
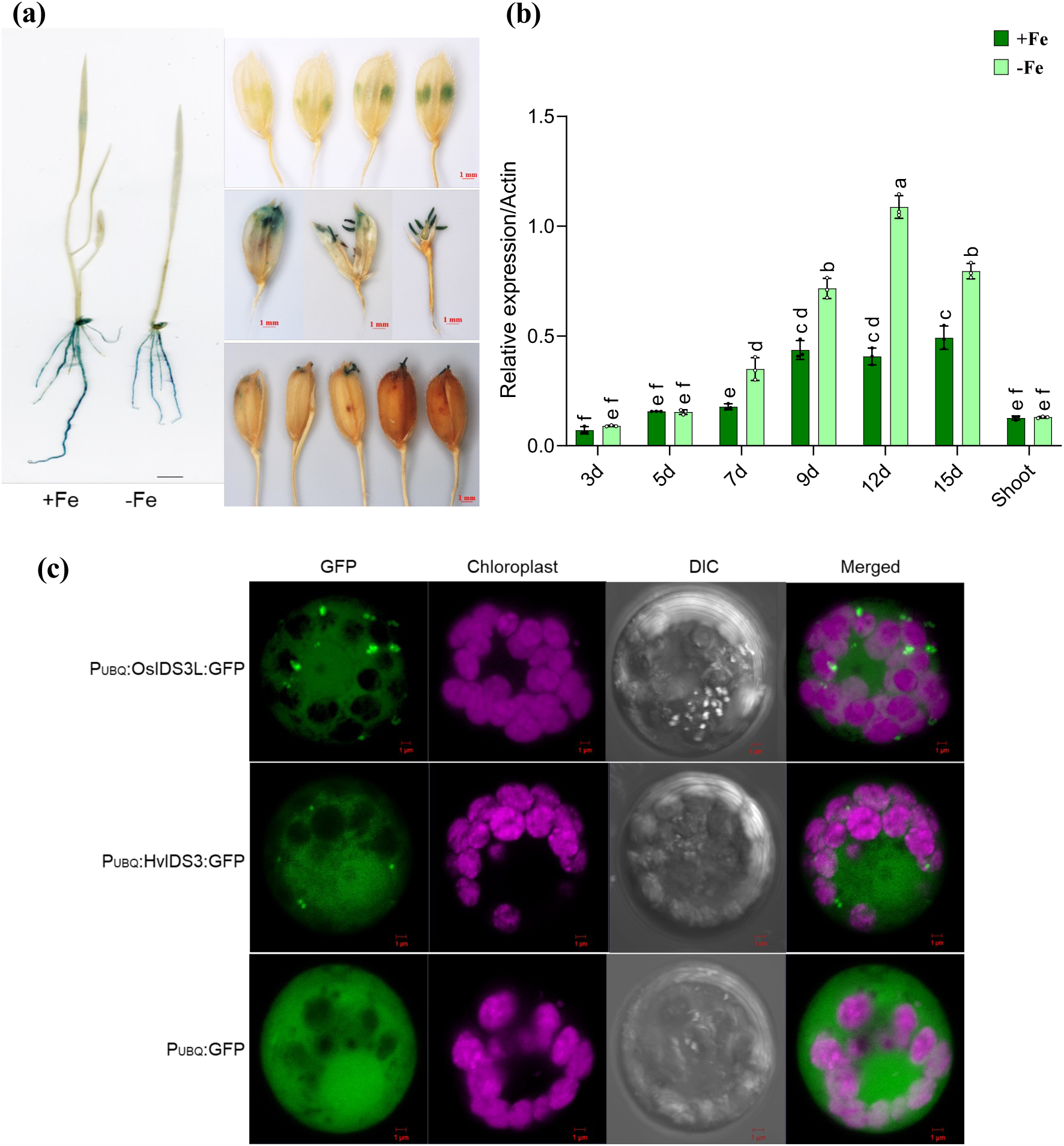
OsIDS3L shows root-specific Fe response and localizes in vesicle-like structures. **(a)** Histochemical GUS staining of P_OsIDS3L_:GUS lines. Left: 10-day-old seedlings grown on ½ MS with and without NaFeEDTA. Intense GUS activity (blue) is observed in roots. Right: GUS staining in a rice flower at different stages shows GUS activity in anthers. **(b)** RT-qPCR expression analysis of OsIDS3L in roots at different time points and 12-day-old shoot samples. Plants were grown on ½ MS medium with or without 50µM NaFeEDTA (+Fe/-Fe) for the indicated number of days. Data represent Mean ± SD, n=3. Different letters indicate significant differences determined by a two-way ANOVA followed by Tukey’s post hoc test. **(c)** Subcellular localization of OsIDS3L in rice protoplast. Confocal images presenting aggregated dot-like structure in wild-type protoplasts transiently expressing GFP fused OsIDS3L and HvIDS3 (P_UBQ_:OsIDS3L:GFP, and P_UBQ_:HvIDS3:GFP). Empty vector (P_UBQ_:GFP) control were used as negative control.

To further quantify Fe-responsive expression, we performed RT-qPCR on a 15-day time course of Fe deficiency. Expression of *OsIDS3L* in roots is induced upon Fe deficiency, with transcript levels increasing progressively up to day 12, followed by a decline at day 15 (Fig. 2b). Besides, *OsIDS3L* transcript abundance under Fe-sufficient conditions suggests OsIDS3L’s significance in general plant growth. Furthermore, minimal expression in 12-day-old shoots suggests that OsIDS3L is root-specific and Fe-responsive. This expression pattern is similar to Fe-responsive genes involved in PS biosynthesis, *OsNAS2* and *OsDMAS1* (Fig. S5).

Since OsIDS3L is predicted as an enzyme, subcellular localization may hint at its enzymatic function. For this purpose, we transiently expressed P*_UBQ_*:*OsIDS3L:GFP* in rice protoplast alongside positive (P*_UBQ_*:HvIDS3:GFP) and negative control (P*_UBQ_*:GFP). *OsIDS3L* and *HvIDS3* transfected protoplasts showed GFP signal aggregated in several dot-like structures (Fig. 2c). These dot-like structures suggest the presence of vesicle-like structures derived from rough endoplasmic reticulum, as reported previously for PS biosynthesizing genes (Nishizawa and Mori, 1987; Nozoye *et al*., 2014). In conclusion, the results suggest that OsIDS3L may be involved in the biosynthesis of PSs and contribute to Fe uptake and homeostasis.

### OsIDS3L overexpression confers tolerance to iron deficiency and enhances yield traits

To investigate the function of OsIDS3L, we generated transgenic rice lines constitutively overexpressing *OsIDS3L.* Three independent lines (OsIDS3L-OE1, OE2, OE3) with confirmed expression were compared to empty vector control (EV) and a positive control (HvIDS3-OE; Hv-OE) (Fig. S6).

Phenotypic analysis was conducted under Fe-deficient conditions induced by (i) Fe deficiency: not adding Fe source in half-strength MS medium, (ii) drought stress induced by using 3% agar, and (iii) alkaline stress induced by increasing pH of the media to 10 (Lindsay and Schwab, 1982; Fan *et al*., 2012; Slessarev *et al*., 2016; Carrijo, Lundy and Linquist, 2017; Calabrese and Porporato, 2019; Kanwar, Baby and Bauer, 2021; Bakaram *et al*., 2022). Results show that OsIDS3L-OE plants have higher shoot length and/or chlorophyll content in all Fe-deficient conditions in comparison to EV control plants (Fig. 3a–c). This suggests enhanced Fe-deficiency tolerance in the OsIDS3L-OE lines. Besides, Hv-OE shows prominent tolerance under alkaline conditions as reported previously (Suzuki *et al*., 2008). However, OsIDS3L-OE plants shows higher tolerance than HvIDS3-OE lines, suggesting *OsIDS3L* is highly efficient in Fe uptake and/or homeostasis.

**Fig. 3.**
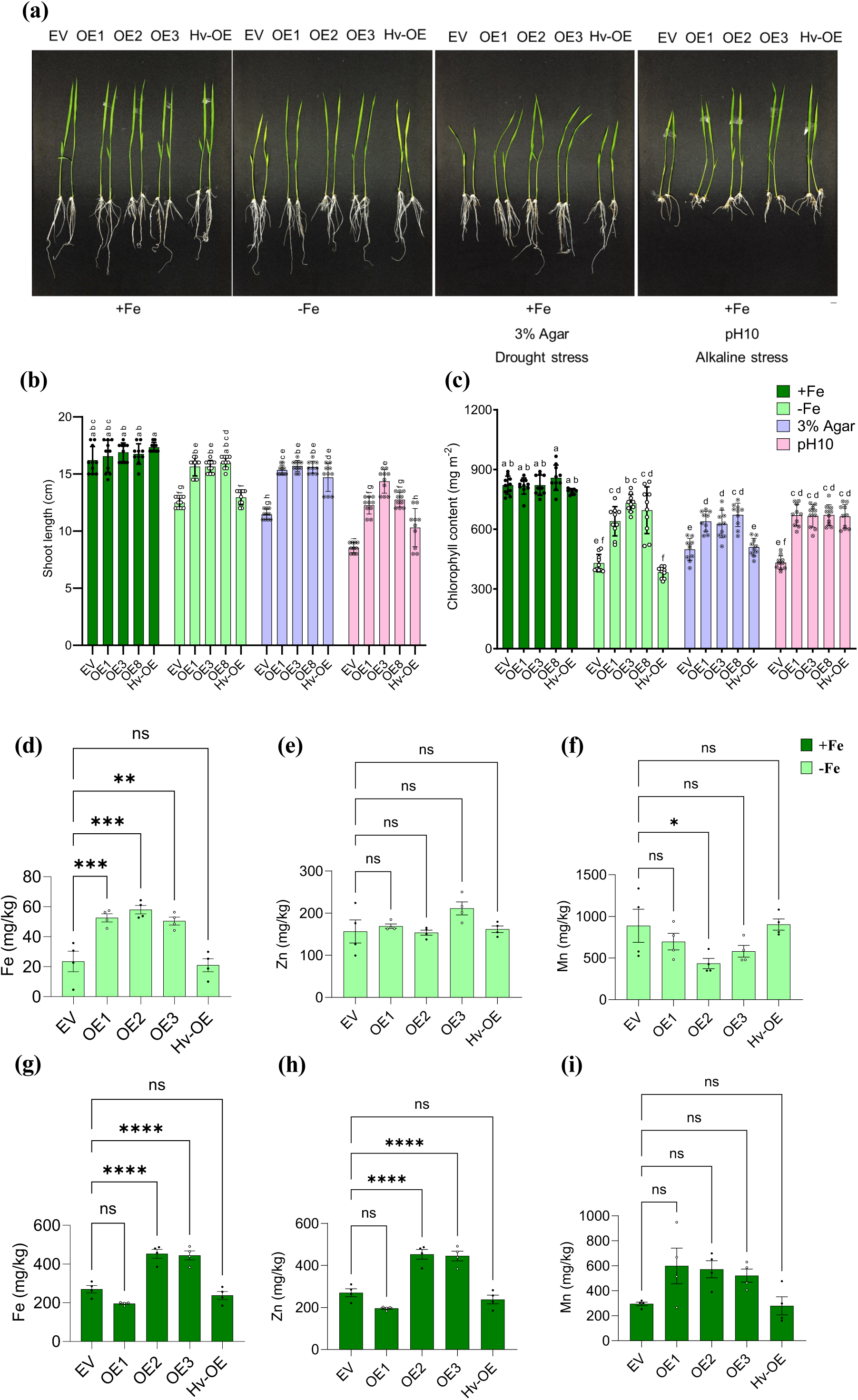
OsIDS3L overexpression confers tolerance to iron deficiency. **(a)** Phenotypic demonstration of 8-day-old OsIDS3L overexpression (OE) rice lines and empty vector (EV) controls under iron-sufficient (control), iron-deficient (no added Fe), 3% agar (drought induced Fe deficiency), and pH10 (alkaline stress induced Fe-deficiency) stress. Scale bar = 1 cm. **(b)** Shoot length and **(c)** chlorophyll content in the youngest leaf of 14-day-old seedlings. Data are presented as mean ± SD (n = 10). Different letters indicate significant differences determined by two-way ANOVA followed by Tukey’s post hoc test. **(d-i)** Micronutrient content (Fe, Zn and Mn) in young leaves of 3-week-old plant. Plants were grown on ½ MS medium with or without 50µM NaFeEDTA. Data represent mean ± SEM, n=4. Statistical analysis: one-way ANOVA followed by Dunnett test, ****P < 0.0001, ***P < 0.0002, **P < 0.0021, *P < 0.0332.

To assess whether the observed tolerance was associated with Fe uptake and homeostasis, we quantified Fe content in young leaves of 3-week-old seedlings using ICP-OES under Fe-deficient and Fe-sufficient conditions. Results show OE lines accumulate nearly 2-fold Fe in comparison to EV under Fe-deficient conditions, explaining Os-OE plant’s better growth (Fig. 3d). Other micronutrients, including Zn and Mn, remain unchanged or slightly affected in Fe-deficient conditions (Fig.3e–f). Interestingly, Fe and Zn levels elevate in two of the OE lines, even under Fe-sufficient conditions (Fig. 3g–i). This suggests that *OsIDS3L* affects micronutrient uptake and homeostasis, which can be beneficial or toxic to the plant, depending on growth conditions.

### OsIDS3L overexpression enhances growth and yield traits in water-saving conditions

Field-related plant growth was determined in alkaline and water-saving conditions. Experiments corresponding to alkaline soil (created by adding calcium oxide) failed as the soil came to equilibrium pH after a month of plant growth, possibly due to soil chemistry as reported earlier (Ponnamperuma, 1972; Hong *et al*., 2018). Therefore, we focused on drought stress. For this purpose, we transferred 8-day-old seedlings to either waterlogging or to alternate watering-drying (AWD, water-saving) conditions. Results show a reduction in panicle number and seed yield in EV control in water-saving conditions (Fig. 4a-c). On the contrary, under water-saving conditions, OE lines yield higher panicle number and seed yield in comparison to EV control (Fig. 4a-c). However, OE shows reduced panicle number and seed yield under waterlogging conditions, which might be the effect of Fe toxicity in the plant as indicated by bronzing phenotype in OE plants (Fig. S9). Furthermore, ICP-OES analysis of seeds shows higher Fe, Zn, and Mn content in OE seeds in comparison to EV control plant seeds (Fig. S7). This suggests that OsIDS3L-OE plants enhance nutrient flow into the seeds, similar to the long-distance micronutrient transport by PSs (Ma and Nomoto, 1996; Connorton and Balk, 2019; Aleksza *et al*., 2024). Conclusively, overexpression of *OsIDS3L* enhances Fe-deficiency tolerance and contributes to micronutrient accumulation in seeds.

**Fig. 4.**
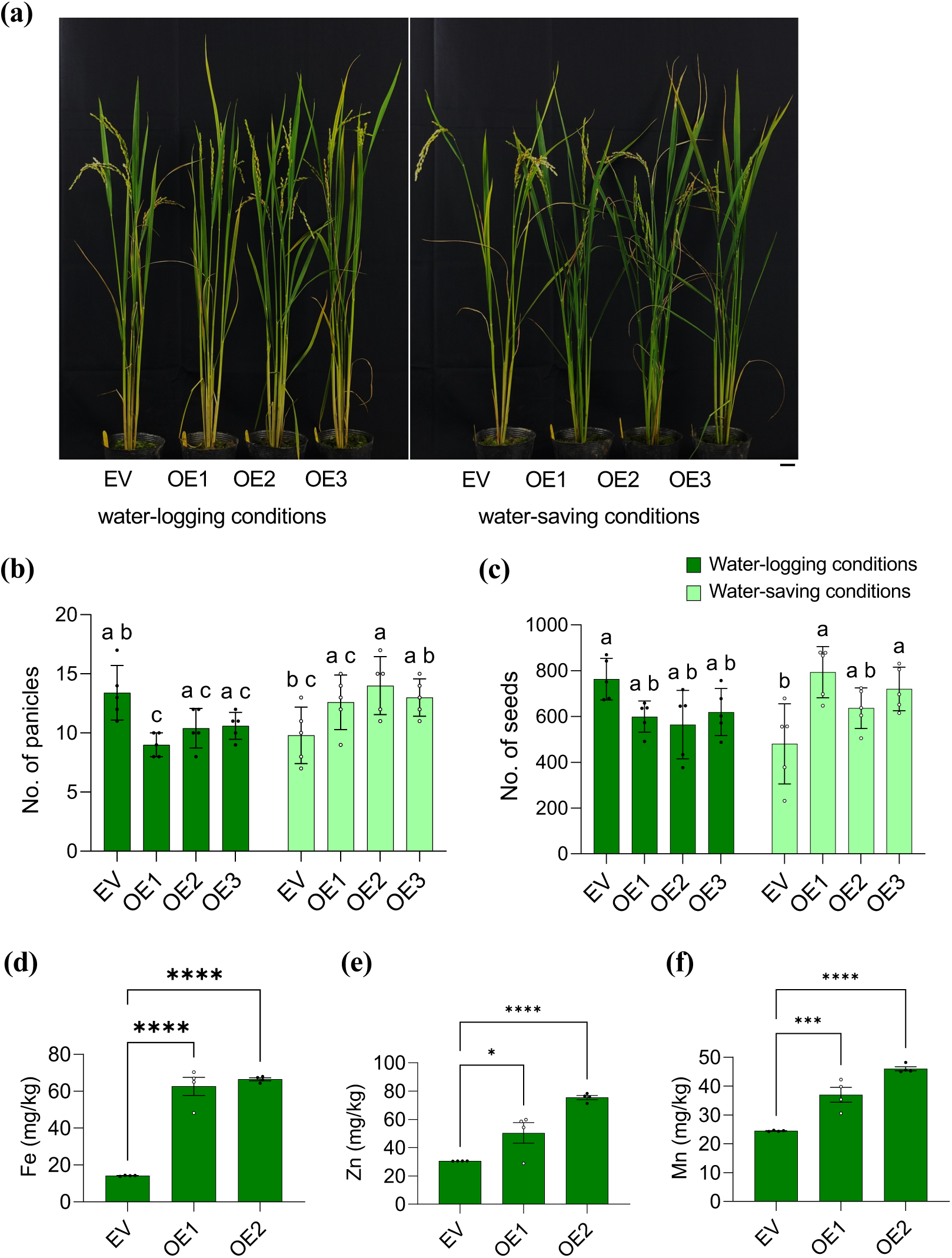
OsIDS3L overexpression lines possess useful agronomic traits. **(a)** Phenotypes of plants grown under water-logging and water-saving cultivation conditions. Scale bar = 10 cm. **(b)** Panicle number, **(c)** seed yield per plant in respective growth condition. Three biological repeats were performed, two in greenhouse and one field experiment. Phenotype images are depicted from greenhouse experiments. Quantitative data is present from field experiment (mean ± SD, n=5). Different letters indicate significant differences as determined by a two-way ANOVA followed by Tukey’s post hoc test. **(d-f)** Micronutrient content (Fe, Zn and Mn) in rice seeds grown performed using ICP-OES. ICP-OES was performed on seeds harvested from water-logging condition. Data shown from 4 biological repeats with 10-15 seeds in each biological repeat. Data represent Mean ± SEM, calculated using one-way ANOVA followed by Dunnett test, ****P < 0.0001, ***P < 0.0002, **P < 0.0021, *P < 0.0332.

### OsIDS3L overexpression alters phytosiderophore levels but does not produce mugineic acid

To investigate the biochemical basis of Fe-deficiency tolerance in OsIDS3L-overexpressing (OE) lines, we conducted LC-ESI-Q-TOF-MS/MS. Since OsIDS3L is homolog of MA-synthesizing enzyme, we looked for MA peak and fragmentation spectra in OsIDS3L-OE roots. However, MA was not detected in OsIDS3L-OE lines (Fig. 5a, S8a), whereas it was readily detected in Hv-OE roots. This indicates that, unlike barley IDS3, OsIDS3L does not synthesize MA in rice. Instead, OsIDS3L overexpression impacted the levels of MA-pathway intermediates, DMA and NA (Fig. 5b-c, S8b-c). NA levels are found to be high in both conditions in OsIDS3L-OE lines. While the DMA levels are higher in OE lines in Fe-sufficient conditions, the levels are similar to EV control lines under Fe-deficient conditions. This suggests that OsIDS3L activity increases the NA and DMA pools in the plant.

**Fig. 5.**
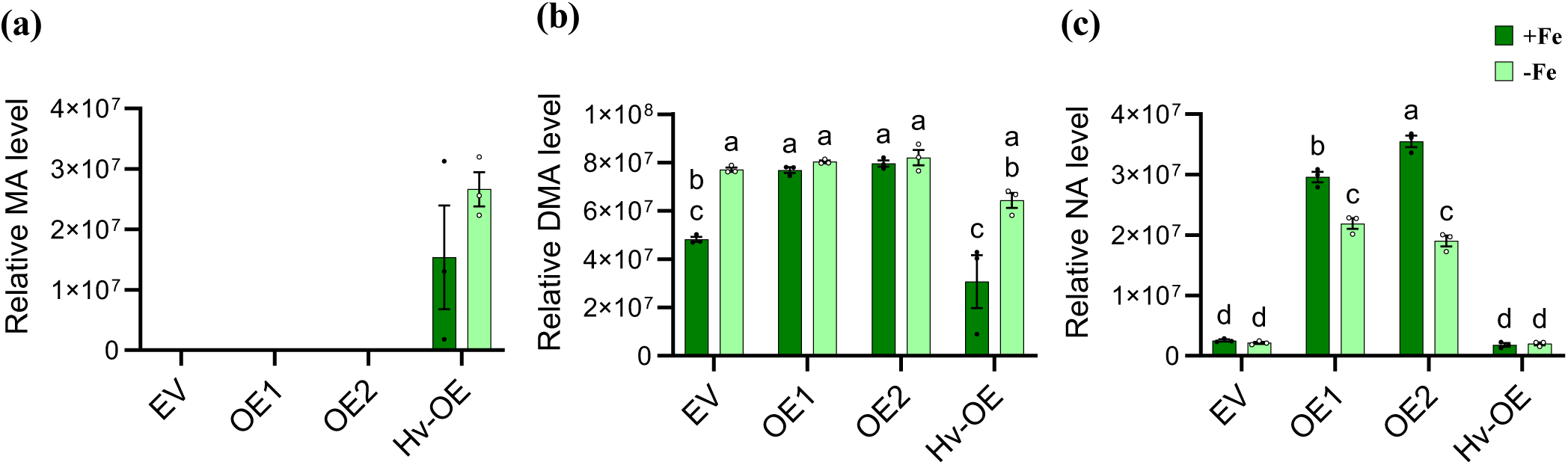
OsIDS3L overexpression (OE) lines accumulate Fe deficiency induced metabolite content. Relative levels (Peak area) of **(a)** mugineic acid (MA), **(b)** deoxymugineic acid (DMA), and **(c)** nicotianamine (NA) contents in 8-day-old root extracts, representing key components of the phytosiderophore (PS) pathway. Plants were grown on ½ MS medium with or without 50µM NaFeEDTA for 8 days. Data are presented as Mean ± SEM, n = 3. Different letters indicate significant differences determined by a two-way ANOVA followed by Tukey’s post hoc test.

### OsIDS3L knockout mutants exhibit normal iron deficiency response as wild type

To further examine OsIDS3L function, we generated knockout (KO) lines using CRISPR-Cas9. Three transgenic lines with no detectable full-length transcript were selected for experiments (Fig. S9). The *dmas1* mutant was also generated using CRISPR as a control to Fe deficiency. Phenotypes comparable to WT were observed for *ids3l* in both Fe-sufficient and deficient conditions (Fig. 6a-c). While *dmas1* displayed strong sensitivity to Fe deficiency. The *ids3l* were further observed in soil. However, it also did not show a distinct phenotype from the wild type. Since OsIDS3L is hypothesized to act downstream of DMA, a subtle or absent phenotype in the KO lines was not unexpected. Furthermore, genes involved in secondary metabolism and adaptive evolution, including other 2OGD members, function in a condition- or tissue-specific manner (Lloyd and Meinke, 2012; Kawai, Ono and Mizutani, 2014; Kinsler, Geiler-Samerotte and Petrov, 2020; Kreutzmann *et al*., 2021; Ament-Velásquez *et al*., 2022; Xu *et al*., 2023). Therefore, absence of visible phenotypes in *ids3l* does not rule out functional activity, and hints to metabolic significance similar to secondary metabolite mutants (Haughn *et al*., 1991; Ishimaru *et al*., 2011; Zhang *et al*., 2013; Ezoe, Shirai and Hanada, 2021; Cullen and Lingwan, 2024). Biochemical analysis of *ids3l* roots shows normal levels of NA and DMA as observed in WT (Fig. 6d-e). On the contrary, substrate accumulation, a higher level of NA was observed in *dmas1* along with unchanged DMA levels as reported previously (Bashir *et al*., 2017). The unchanged DMA level in *ids3l* hints at (i) an alternate enzyme with a similar function, (ii) an alternate substrate, or (iii) metabolic redundancy from a parallel pathway. However, OsIDS3L is a single-copy gene; therefore, the remaining possibilities include alternate substrates and metabolic redundancy.

**Fig. 6.**
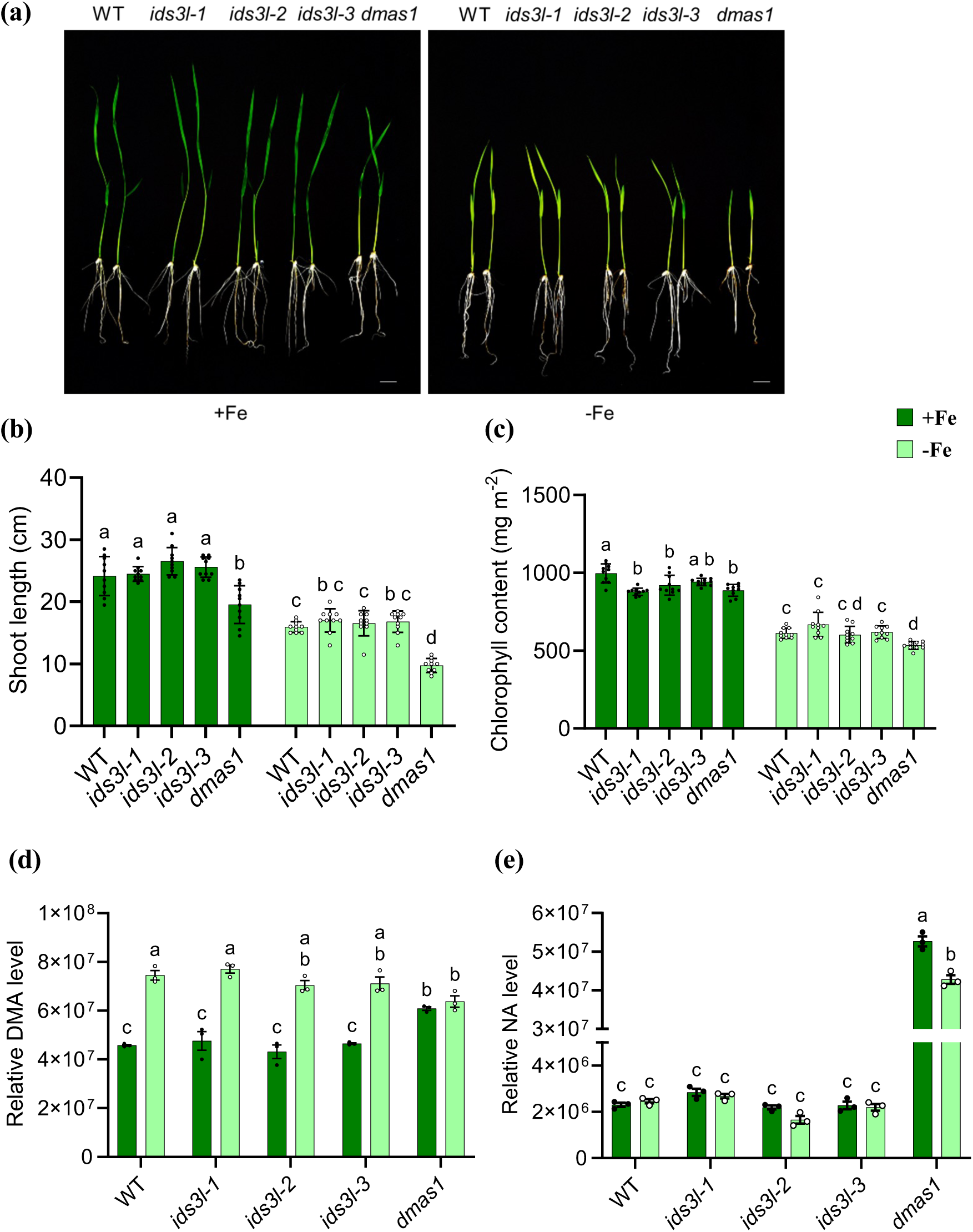
OsIDS3L knockout mutants exhibit similar iron deficiency responses as the wild type (WT). **(a)** Phenotypic demonstration of WT, *ids3l*, and *dmas1* seedlings grown on ½ MS medium with or without 50µM NaFeEDTA for 8 days. Scale bar = 1cm. **(b)** Shoot length of and **(c)** chlorophyll content in the youngest leaves from 3-week-old seedlings germinated and grown on ½ MS medium with or without iron. Data are presented as mean ± SD, n = 10. **(d)** Deoxymugineic acid (DMA) and **(e)** nicotianamine (NA) relative levels (peak area) in 8-day old root extracts from plants germinated and grown on ½ MS medium with or without 50µM NaFeEDTA. Data represent mean ± SEM, n = 3. Different letters indicate significant differences determined by a two-way ANOVA followed by Tukey’s post hoc test.

### Untargeted metabolomics suggests an OsIDS3L-dependent pathway in phytosiderophore metabolism

To investigate the potential enzymatic function of OsIDS3L, we profiled the root metabolome of WT, KO, and OE. PCA of metabolic features shows that 34% of metabolic variance (PC1) is affected by Fe status, and 14% (PC2) affected by genotypes. Compact and non-overlapping confidence ellipses (95%) shows strong reproducibility. PERMANOVA analysis further supports these findings (F = 35.4, R² = 0.94, p = 0.001) (Fig. S10a). Pairwise PERMANOVA contrasts (all p < 0.05) suggest that minor ellipse overlap among PCA groups is attributed to shared Fe status. Hierarchical clustering based on Euclidean distance corroborates the PCA analysis (Fig. S10b). In conclusion, OE and KO samples cluster closely, distinct from WT in the control condition, suggesting that both gain/ loss-of-function of OsIDS3L affect baseline metabolism (Fig. S10a-b). Furthermore, under Fe-deficient treated (T) conditions, OE forms a distinct cluster, consistent with its Fe-deficiency-tolerant phenotype (Fig. S10a-b). A heatmap of the 100 most variable features (Fig. S10c) revealed distinct metabolic clusters. Cluster A (Fig. S10c) suggests Fe-responsive metabolite features. These OsIDS3L unaffected features suggest particular role of OsIDS3L in metabolism. Cluster B and C (Fig. S10c) suggests features whose levels differed by genotype irrespective of condition. This suggests OsIDS3L alters certain metabolic pathways constitutively.

To pinpoint metabolites directly related to OsIDS3L’s enzymatic activity, we employed feature-based molecular networking (FBMN). The molecular network groups metabolites by similarity (cosine score) in MS/MS fragmentation spectra (indicating structural relatedness). Since OsIDS3L is a dioxygenase enzyme, it is expected to catalyze a hydroxylation reaction. Therefore, edges with Δm/z ∼ 15.9949 Da; tolerance ±0.02 Da were selected. A total of 49 such oxidative transformation edges were identified (Table S2a). The resulting feature list is targeted to substrate-product levels logic (chemotype), where the putative “product” is more abundant in OE and the “substrate” is more abundant in KO (relative to WT), reasoning that OsIDS3L overexpression should drive product formation, whereas its absence would cause substrate accumulation.

For OE lines, features accumulating at least 1.5-fold higher than WT were designated as potential products. This yielded 19 OE-specific oxidative reactions (Table S2b). In contrast, KO lines did not yield any features at the 1.5-fold threshold, likely due to pathway redundancy or compensatory mechanisms. We therefore relaxed the criterion to 1.2-fold higher in KO lines as potential substrates, yielding 13 KO-specific oxidative reactions (Table S2c). Overlapping OE- and KO-specific oxidative reactions revealed seven candidate reactions (Table S2d). Mapping the seven candidate reactions to the FBMN network revealed a reaction (9686_259.1292_2.49 → 8219_275.1241_2.21, depicted as feature number_m/z_RT) sharing MS/MS fragmentation spectra similarity with DMA (Fig. 7a). Corresponding to the analysis, substrate 9686 is high in KO though it didn’t pass significant test and product, 8219 is higher in OE (Fig. 7b-c, S11).

**Fig. 7.**
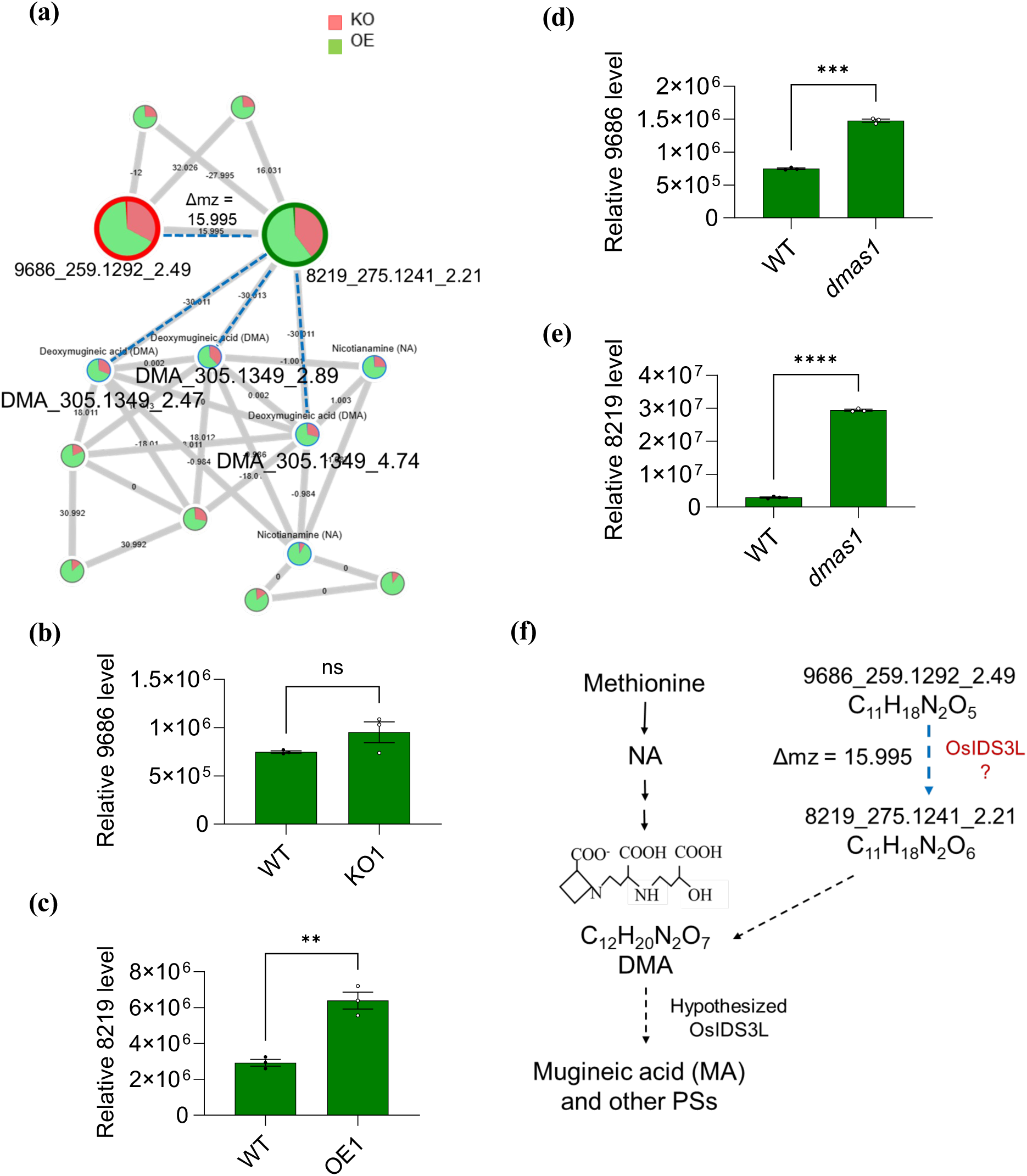
OsIDS3L may catalyze a reaction upstream of DMA. **(a)** GNPS-FBMN network illustrates metabolic features sharing MS/MS fragmentation spectra with DMA. Each node in the network represent a metabolic feature connected by an edge that represent MS/MS fragmentation similarity. The highlighted big nodes, circled in red (9686_259.1292_2.49) is expected substrate and circled in green (8219_275.1241_2.21) is expected product of OsIDS3L enzyme reaction. The edge between the two features shows m/z difference of ∼16 Da which represent hydroxylation reaction. Feature, 8219 share edge with DMA sharing Δm/z of 30 Da. The edges of significance are highlighted in dashed blue lines. The concentration of metabolites in OE and KO is presented by shaded green and red area in nodes. **(b-e)** Relative level of OsIDS3L proposed enzyme’s substrate and product. **(b)** substrate 9686 in KO (expected to increase), **(c)** product 8219 in OE (expected to increase), **(d)** 9686 in *dmas1,* **(e)** 8219 in *dmas1.* Data represent mean ± SEM, n = 3. Statistical analysis: Welsch’ t test, ****P < 0.0001, ***P < 0.0002, **P < 0.0021, *P < 0.0332. **(f)** Proposed pathway for OsIDS3L enzyme activity. The hypothesized OsIDS3L enzyme activity was defined from DMA to MA like HvIDS3. Based upon the computational analysis of untargeted metabolomics, the new proposed OsIDS3L enzyme activity is indicated in blue dashed arrows and red enzyme label catalyzing a hydroxylation reaction upstream or parallel to DMA biosynthesis pathway.

The FBMN network shows three features corresponding to DMA (Fig. 7a), two of which correspond to the same feature, distinguished by a broader peak (Retention time-RT), and the third hints at a stereoisomer based on high-confidence spectral similarity (Fig. S12). The edges connecting 8219_275.1241_2.21 to DMA features show a Δm/z ≈ 30 Da, which resonates with a CH₂O (formaldehyde) group as indicated by Modifinder (Fig. S13). In-silico structure prediction tool, SIRIUS predicted molecular formulae of 9686_259.1292_2.49 as C₁₁H₁₈N₂O₅ and 8219_275.1241_2.21 as C₁₁H₁₈N₂O₆, presenting a difference of a single oxygen molecule between substrate and product. Furthermore, the molecular formulae for feature 8219_275.1241_2.21 align with Modifinder’s predicted CH₂O-modification in DMA (C₁₂H₂₀N₂O₇). The results suggest that OsIDS3L may act upstream of DMA, catalyzing a hydroxylation reaction and synthesizing the 8219_275.1241_2.21 feature, which might be formylated/hydroxymethylated (CH₂O addition) to form DMA. This indicates that OsIDS3L may contribute to an alternate biosynthetic pathway intersecting the canonical DMA biosynthesis pathway. This pathway intersection could also explain two observations: (1) the elevated DMA levels in *OsIDS3L*-OE roots, and (2) unaffected DMA levels in *dmas1* mutants. We analyzed the *dmas1* mutant for 9686_259.1292_2.49 and 8219_275.1241_2.21 features, which indicate accumulation of these compounds in *dmas1*, supporting DMA synthesis from an alternate pathway (Fig. S14). Together, these data suggest that OsIDS3L may catalyzes an oxidation reaction branched upstream of the DMA pathway and contributes to Fe homeostasis through a non-canonical DMA biosynthesis pathway (Fig. 7f).

### OsIDS3L overexpression reprograms iron-deficiency responses in roots

While the untargeted metabolite analysis suggests an uncanonical DMA biosynthesis pathway, the elevated NA levels in the overexpression (OE) lines remain intriguing. We hypothesize that OsIDS3L OE constitutively activates iron-deficiency responses. To examine transcriptional regulation in gain- and loss-of-function backgrounds, we performed RNA-seq and compared the transcript abundance of Fe-deficiency–regulated genes.

The OE lines showed markedly higher expression of Fe-regulated genes under control conditions, whereas these same genes displayed attenuated induction under Fe-deficiency stress compared with WT (Table 1, Fig. S15a). This pattern indicates a constitutively elevated “iron-deficiency–like” transcriptional state in OE roots, enabling plants to better tolerate Fe-deficient conditions with only minimal additional induction upon stress (Table 1, Fig. S15).

**Table 1.**
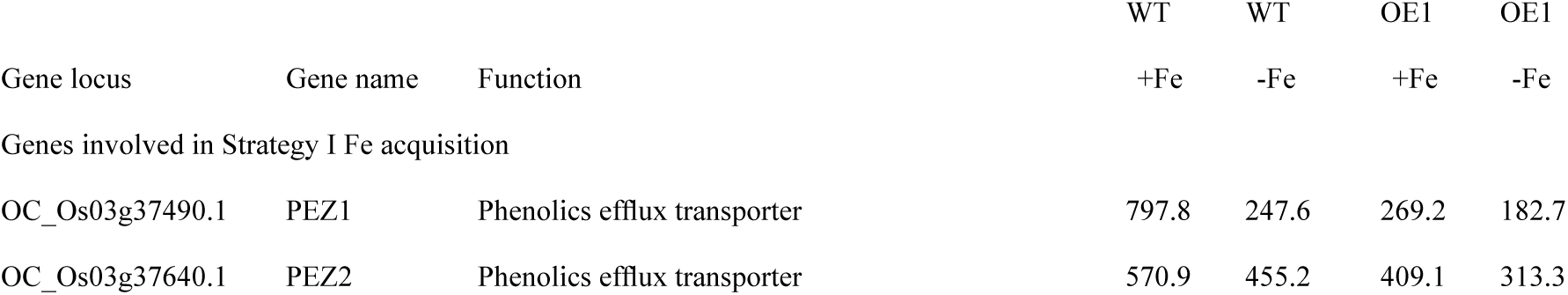
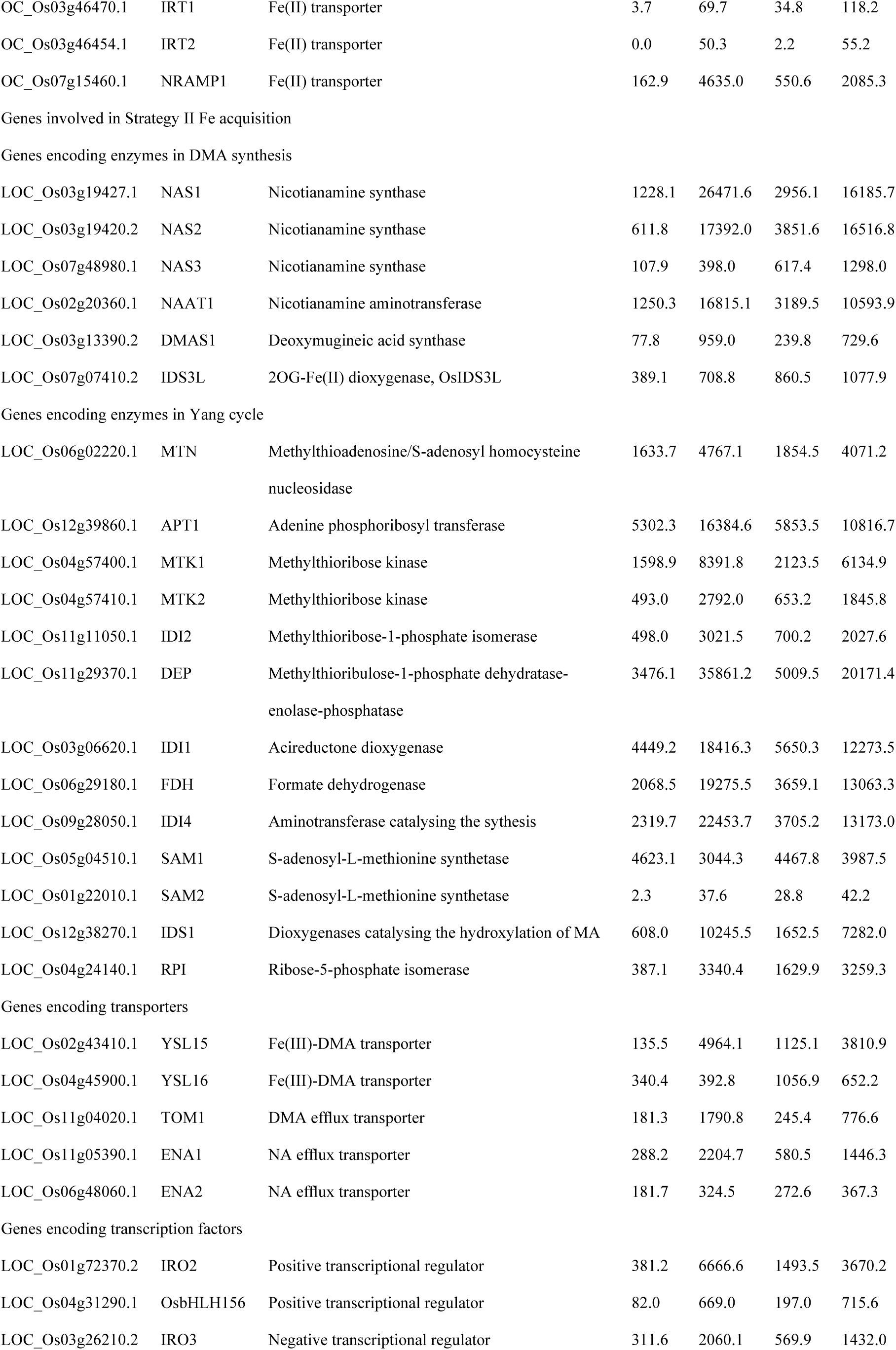
DESeq2 values for genes involved in Strategy I and II Fe acquisition in roots of wild-type (WT) and OsIDS3L overexpression (OE1) lines. The DESeq2 values were obtained from average of three biological repeats. Gene list adopted from (Wang *et al*., 2020).

In contrast, the KO lines exhibited only mild changes in the expression of Fe-deficiency–regulated genes relative to WT (Table S3), and the overall expression pattern largely mirrored that of WT. This limited impact is consistent with common observations that loss of a single enzyme in a branched or compensated metabolic network often results in minimal transcriptional changes, because parallel pathways, functional redundancy, or homeostatic regulation can buffer against the loss of function. Thus, transcriptional regulation in KO roots appears much less perturbed than in OE roots. Taken together, the increased NA and DMA accumulation is consistent with the transcriptional reprogramming observed in the OE lines.

## Discussion

### OsIDS3L, HvIDS3 homolog with functional divergence from HvIDS3

In this study, we identified OsIDS3L, a Fe(II)/2-oxoglutarate-dependent dioxygenase (2OGD) enzyme homologous to HvIDS3 (Fig. 1). The OsIDS3L enzyme shares a 60% homolog similarity which is quite reasonable to study the enzyme, given that the percentage similarity between HvIDS2 and HvIDS3 is just 54% (Fig. 1c). Being a single-copy gene in rice (Fig. S2-3), it hints to an essential role in rice. The Fe deficiency response of OsIDS3L in roots indicates its role in Fe uptake (Fig. 2a-b). Furthermore, GUS staining was observed in anthers, that indicates OsIDS3L’s role in Fe homeostasis and development, possibly nutrient supply (Fig. 2a). This could result in tolerance to Fe deficiency of OsIDS3L overexpression (OE) lines in a Fe-deficient medium (Fig. 3), where the stored Fe in rice seeds may facilitate early growth. This tolerance is further extended to later stages of rice growth as observed with higher panicle number and seed yield in OE lines under water-saving cultivation conditions (Fig. 4). These phenotypic observations aligned with the predicted function of the enzyme, that is, enhanced Fe uptake and homeostasis by mugineic acid (MA) biosynthesis.

Surprisingly, the metabolic analysis revealed no signs of MA in OE lines which can be detected in HvIDS3-overexpressing (Hv-OE) lines used as a positive control (Fig. 5). Besides MA synthesis, our phenotypic analysis shows Hv-OE line cannot cope Fe deficiency stress except high pH (observed phenotype in earlier report (Suzuki *et al*., 2008)) (Fig. 3). Previous report suggests that constitutive expression of HvIDS3 cDNA was less efficient than the 20kb genome fragment containing HvIDS3 promoter (Kobayashi *et al*., 2001). Therefore, it is possible that HvIDS3 and MA synthesis/quantity (PS quantity is an important trait in Fe deficiency tolerance (Hiradate, Ma and Matsumoto, 2007; Tsednee *et al*., 2012)) is regulated within rice. Conclusively, when using coding sequence, the rice homolog, OsIDS3L has a role in Fe uptake and homeostasis, more efficient than HvIDS3’s MA. The lack of MA in OE lines also suggests synthesis of a different product than MA indicating functional divergence (neofunctionalization) of IDS in rice. We further investigated the predicted substrate (deoxymugineic acid, DMA) concentration. The overexpression may result in a reduction in substrate, as overexpression of the enzyme will readily convert it to the product. The metabolic analysis depicted higher levels of DMA and nicotianamine (NA, precursor of DMA) (Fig. 5). These results were against our hypothesis of predicted substrate. However, the enzyme demand for substrate and its metabolic flux could explain the higher level of pathway compounds. However, this enhanced DMA and NA could explain the Fe deficiency tolerance phenotype along with micronutrient accumulation in seeds. DMA and NA are well known for Fe uptake and homeostasis within the plant, including low-affinity uptake of other micronutrients like Zn and Mn (Ma and Nomoto, 1996; Aleksza *et al*., 2024). Quantitative enhancement of DMA and NA, therefore, corresponds to enhanced Fe uptake and homeostasis as reported earlier (Hiradate, Ma and Matsumoto, 2007; Tsednee *et al*., 2012).

### OsIDS3L enzymatic activity is auxiliary but not essential

To understand the substrate specificity and loss-of-function effect of OsIDS3L, we studied the mutant. As expected, *ids3l* knockout (KO) lines displayed no overt phenotypes under the tested conditions (Fig. 6), since DMA synthesis is sufficient for the plant to overcome Fe deficiency stress. Although the KO mutant shows no phenotype under standard conditions, this is common in secondary metabolism that acts under severe stress or that has unknown indirect compensatory mechanisms. This is also consistent with the physiology of genes involved in secondary metabolism resulting in adaptive evolution, where phenotypic effects are often subtle or conditional (Lloyd and Meinke, 2012). Furthermore, metabolic analysis of KO lines showed no reduction in DMA levels (Fig. 6). In the complex metabolic networks, it is hard that metabolic content gets affected (Alseekh and Fernie, 2018). However, this prompted us to reconsider DMA as the substrate of OsIDS3L. To determine the role of OsIDS3L in DMA to MA synthesis, we also attempted in-vitro enzyme reactions by generating recombinant protein. However, we failed even with positive control, HvIDS3. This is perhaps because of the enzyme contradictory aspect where the dioxygenase require Fe(II) and oxygen for enzyme reaction, which in-vitro is converted to Fe(III) and chelated by the substrate, DMA as mentioned in earlier reports(Nakanishi *et al*., 1993, 1997, 2000; Kobayashi *et al*., 2001).

### OsIDS3l may act upstream of DMA ensuring DMA synthesis via an alternate pathway

To identify the enzyme catalyzed metabolic reaction, metabolic features were grouped based on MS/MS fragmentation spectra using the computational tool Feature-Based Molecular Networking (FBMN). The network was generated by removing noise from blank data and also removing features not observed in 20% of the samples. Thereby, significant edges corresponding to the hydroxylation reaction difference of ∼16 Da m/z were analyzed. Since complex metabolic networks secure metabolic content from significant effects (Alseekh and Fernie, 2018), a relaxed metabolic phenotype/ chemotype is implicated. Metabolic phenotype/chemotype used to identify the enzyme reaction is that the product accumulates in OE and the substrate in KO. The analysis highlighted a hydroxylation reaction (9686_259.1292_2.49 → 8219_275.1241_2.21, depicted as feature number_m/z_RT) sharing an edge with DMA in the FBMN network (Fig. 7) with an m/z difference of ∼30 Da. The result suggests that OsIDS3L may catalyze a reaction upstream of DMA, where 9686_259.1292_2.49 could be catalyzed by OsIDS3L to 8219_275.1241_2.21. The product of OsIDS3L, 8219_275.1241_2.21 may then undergo a formylation/ demethylation reaction by one carbon assimilating enzyme, perhaps by an unknown formaldehyde-condensation enzyme. This suggests that OsIDS3L may catalyze a reaction upstream of, or parallel to, canonical DMA biosynthesis. The proposed hydroxylation reaction by OsIDS3L can explain high DMA levels in OsIDS3L-OE lines. While NA content is much higher than DMA, the 8219_275.1241_2.21 MS/MS spectral similarity to NA is less than that of DMA (Fig. S14), and therefore, we propose that the reaction is upstream of DMA than NA. Furthermore, just like the increased level of NA in dmas1, the concentration of 9686_259.1292_2.49 and 8219_275.1241_2.21 is high in *dmas1,* which somehow explains the unaffected DMA levels in *dmas1* mutants and supports the OsIDS3L predicted enzymatic reaction. The unchanged DMA level was also observed in reported *dmas1*, and the gene redundancy was determined as one of the possible reasons (Bashir *et al*., 2017). The untargeted metabolomics analysis suggests the role of OsIDS3L in an enzyme reaction upstream of DMA (Fig. 7), which could be the result of neofunctionalization of the enzyme in rice. Although further biochemical validation is required to validate OsIDS3L function. Our study suggests the role of OsIDS3L in the non-canonical phytosiderophore biosynthesis pathway. In plants, such metabolic redundancy from different pathways secures key metabolites and masks the effects of gene loss under standard growth conditions (Lorence *et al*., 2004; Tiski *et al*., 2011; Kasahara, 2016), explaining the lack of an obvious phenotype in *ids3l*.

### OsIDS3L overexpression tolerance is correlated with the induced expression of Fe deficiency-responsive genes

The high NA and DMA content in OE, and unaffected levels in KO, make us question if it is a transcriptional response or metabolic flux. To answer the question, we performed whole root transcriptome analysis. The results suggest constitutive expression of Fe-responsive genes in OE lines (Table 1, Fig. S15). As a result, NA and DMA levels are observed to be high in OE lines even under control conditions (Fig. 5). This might prepare OE lines for Fe-deficiency stress as the expression level of Fe-responsive genes is reduced in OE in comparison to WT control under Fe deficiency (Table 1, Fig. S15). On the other hand, OsIDS3L mutation shows somewhat similar regulation of Fe deficiency responsive genes as WT, correlating with the phenotype and metabolic response (Fig. S3). The analysis indicates that the induced NA and DMA levels in OE could be a response to transcriptional regulation.

Our findings related to OsIDS3L highlight the physiological, evolutionary, and biochemical significance of OsIDS3L in Fe uptake, homeostasis and adaptation to Fe-limiting conditions in rice cultivation. As Fe(II)-2OG dioxygenase enzymes are widely implicated in stress-responsive and adaptive metabolite biosynthesis (Kawai, Ono and Mizutani, 2014; Xu *et al*., 2023). OsIDS3L is likely expected to contribute to evolutionary strategies that enhance plant survival under nutrient stress. As a result, OsIDS3L overexpression confers dual benefits, increased tolerance to Fe deficiency and elevated micronutrient accumulation in seeds. Further, it plays a significant role in water-saving cultivation conditions and stress tolerance. These traits highlight its potential utility in breeding and biofortification programs to improve the nutritional quality and stress resilience of staple crops. While these traits are linked to the distinct metabolism of OsIDS3L-overexpression lines, only 10-12% known metabolites make the study challenging (Shahneh *et al*., 2024). Metabolic networks are usually complex, which often makes it hard for metabolic content to get affected (Alseekh and Fernie, 2018). Further, detecting a possible substrate and product among unknowns is complex. In the manuscript, we could only hint at the function of the OsIDS3L enzyme. Further study of the OsIDS3L enzyme may provide deeper insight into the complex metabolites and metabolic networks that govern Fe uptake and distribution. In conclusion, our manuscript demonstrates that rice harbors an MA-synthesizing homolog that has been overlooked for years. Further, it can help plants tolerate Fe deficiency and accumulate micronutrients in the seeds.

## Acknowledgements

This research was supported by the Ministry of Science and Technology (MOST 110-2313-B-001-008-MY3) and Academia Sinica of Taiwan. We thank the technical support from the Academia Sinica Advanced Optics Microscope Core Facility, the Metabolomics Core Facility of the Agricultural Biotechnology Research Center, the Plant Tech Core Facility, and I-Chien Tang for performing element analysis. We thank Professor Chang-Sheng Wang for the help with field experiments in the transgenic paddy field of National Chung-Hsing University. We thank Dr. Mingxun Wang (University of California, Riverside) for support during GNPS2 Office hours and for suggestions on the analysis of untargeted metabolomics data.

