## Supplementary material for "OsIDS3L, a Non-Canonical Dioxygenase Enhancing Iron Homeostasis and Micronutrient Biofortification in Rice": SI Figures

Authors: **Jyoti Aggarwal**[1,2,3] and **Kuo-Chen Yeh**[1,2,4,*] [1]Agricultural Biotechnology Research Center, Academia Sinica, Taipei 11529, Taiwan [2]Molecular and Biological Agricultural Sciences Program, Taiwan International Graduate Program, Academia Sinica and National Chung Hsing University, Taipei 11529, Taiwan [3]Graduate Institute of Biotechnology, National Chung Hsing University, Taichung 40227, Taiwan [4]Biotechnology Center, National Chung Hsing University, Taichung 40227, Taiwan

The following Supporting Information is available for this article:

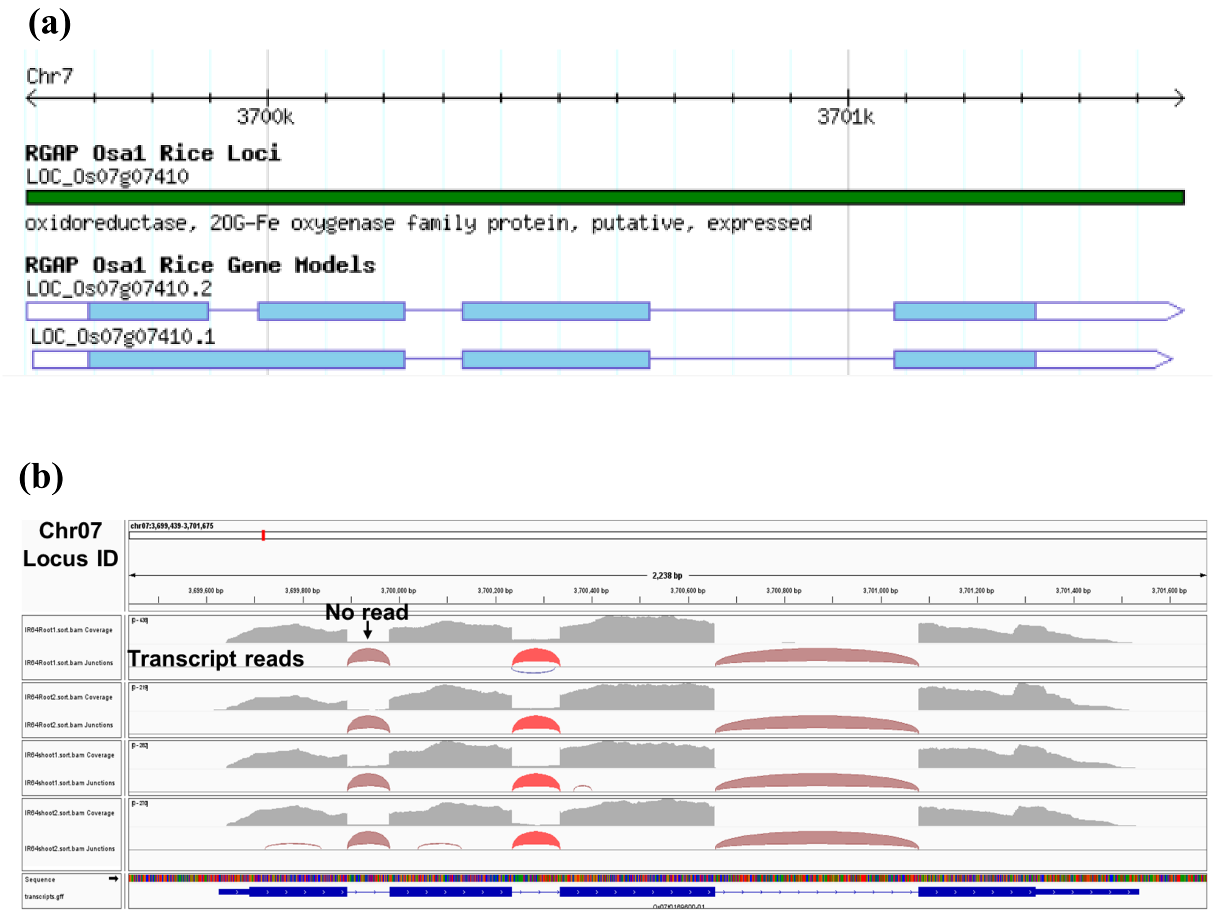


**Fig. S1** Os07g07410.2 is a major transcript. **(a)** Visual representation of splice variants of Os07g07410 from Rice Genome Annotation Project (RGAP). **(b)** RNA-seq data visualization using Integrative Genomics Viewer (IGV) software over Os07g07410 shows no transcript read over first intron. The figure shows the chromosome location in bp at the top. Below it, the grey region shows RNA-seq transcript reads from 4 samples. The reads are missing at three intervals which is intron region. The lowest panel shows the OsIDS3L transcript structure over DNA in blue.

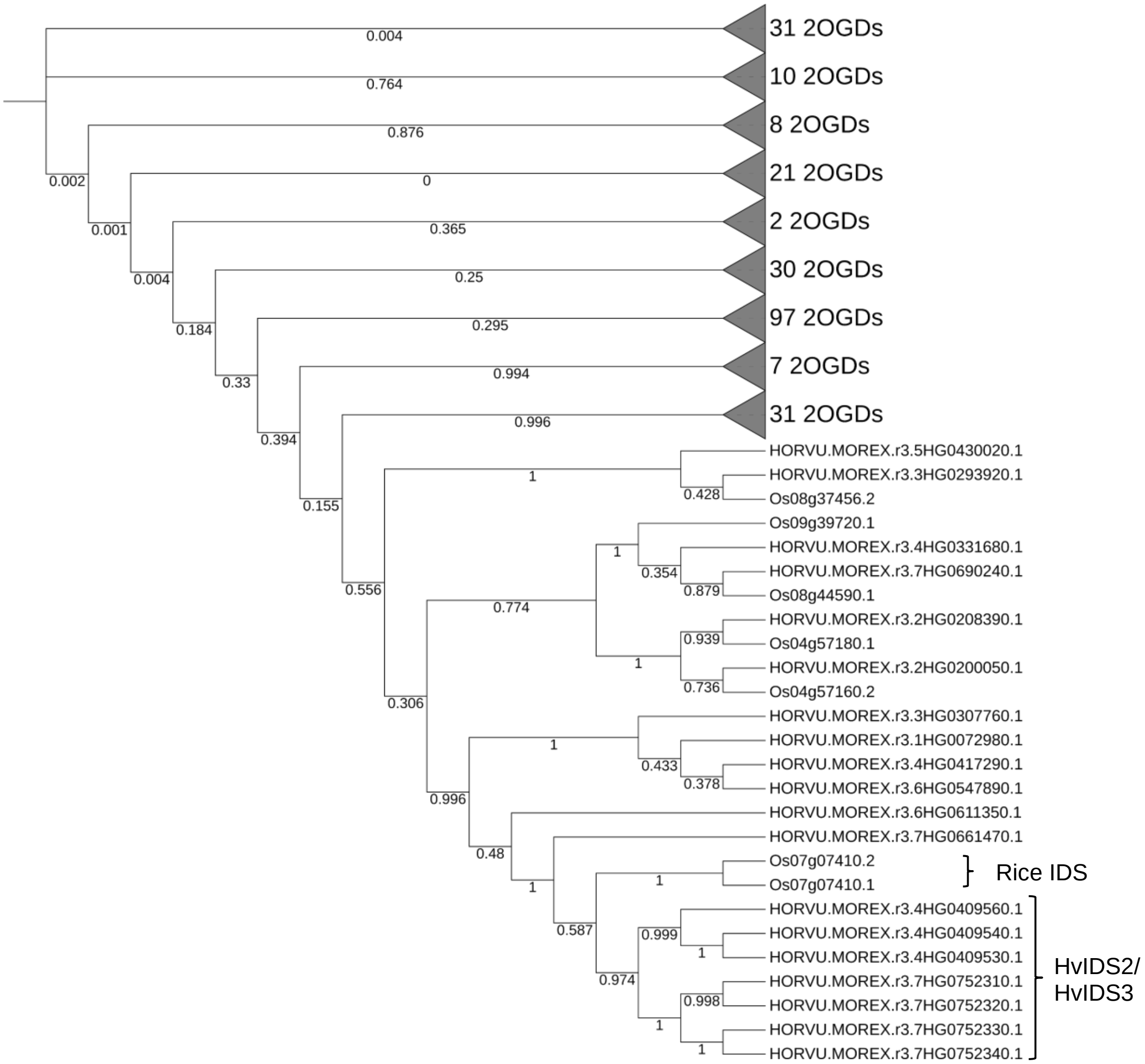


**Fig. S2** Os07g07410 is a single copy gene in rice with homology to barley IDS2/3. Maximum-likelihood phylogenetic tree of Fe(II)-2-oxoglutarate-dependent dioxygenases (2OGDs) from rice and barley with 1000 bootstrap replicates. Rice IDS clusters in the same clade as barley IDS2/IDS3 indicating a common ancestry.

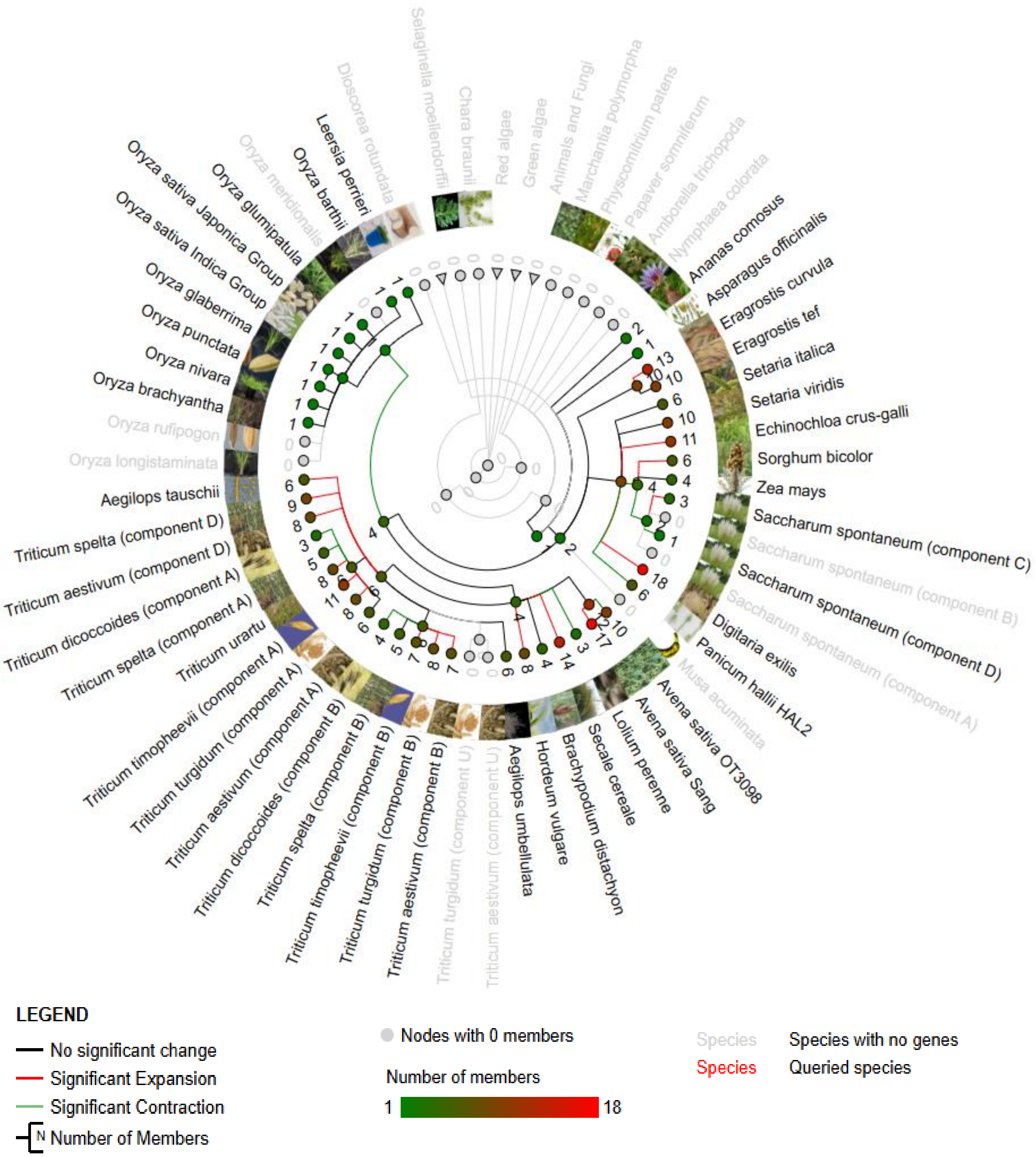


**Fig. S3** Os07g07410 is a single copy gene. The global phylogenetic tree retrieved from EnsemblPlants using Ensembl gene ID; Os07g0169600, EPlGT00940000169430**.** The phylogenetic tree shows the presence of OsIDS3L (one gene) in the tree indicated by the numerical presented at the nodes. The copy is conserved among graminaceous plants and absent in other (zero copy in the species in grey). The species in grey with zero copy also includes *Oryza rufipogon, Oryza longistaminata and Oryza meridionalis*. However, the homolog search shows the conserved gene copy which is unannotated in these genomes.

**(a)**

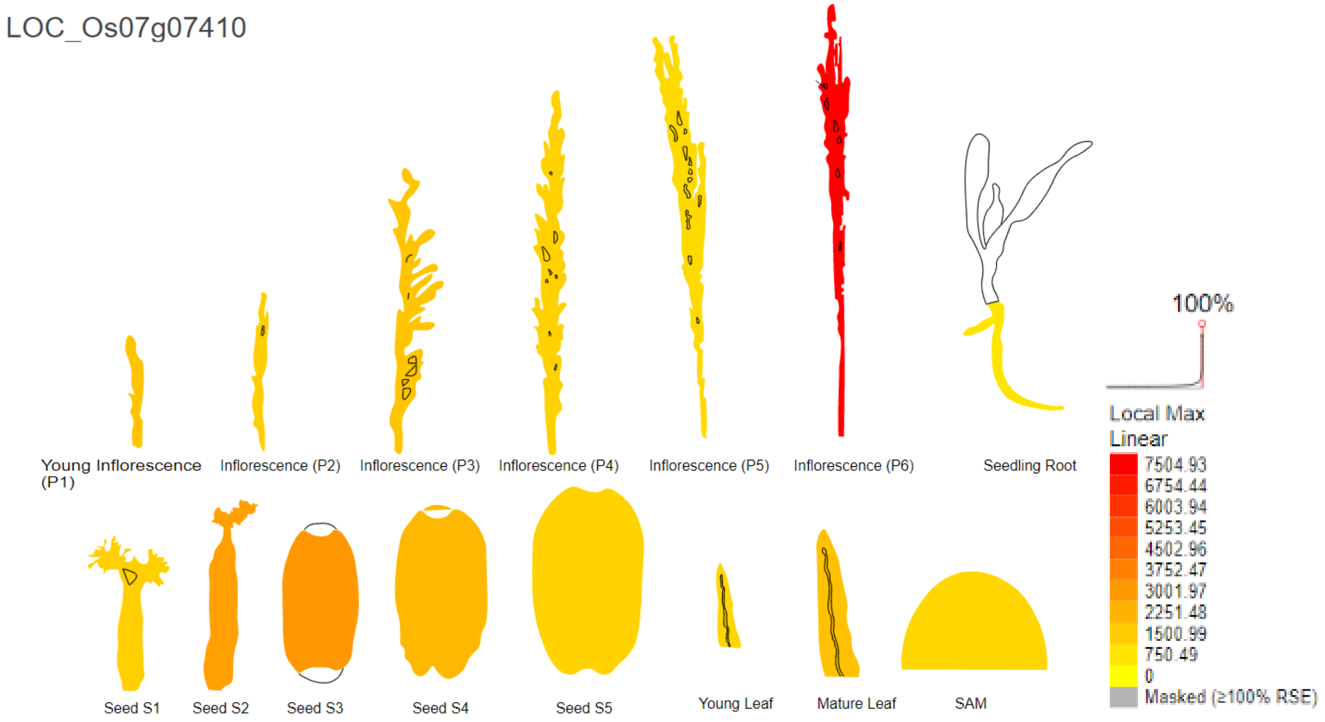
LOC_Os07g07410
Young Inflorescence (P1)
Inflorescence (P2)
Inflorescence (P3)
Inflorescence (P4)
Inflorescence (P5)
Inflorescence (P6)
Seedling Root
Seed S1
Seed S2
Seed S3
Seed S4
Seed S5
Young Leaf
Mature Leaf
SAM
100%
Local Max
Linear
7504.93
6754.44
6003.94
5253.45
4502.96
3752.47
3001.97
2251.48
1500.99
750.49
0
Masked (≥100% RSE)

**(b)**

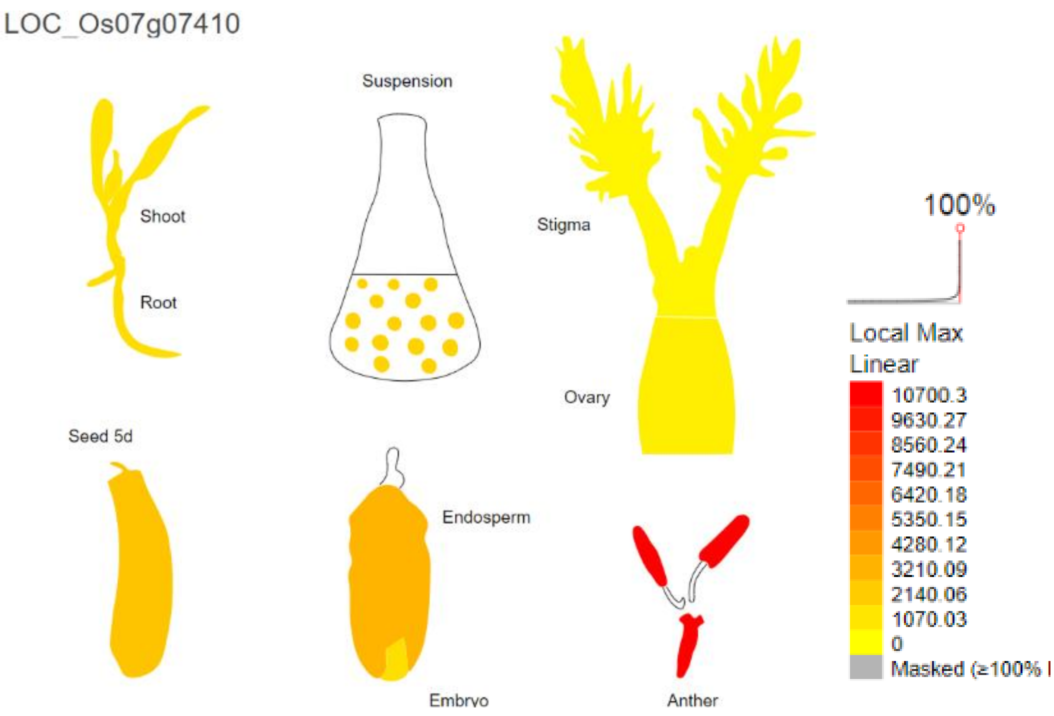
LOC_Os07g07410
Shoot
Root
Suspension
Stigma
Ovary
Seed 5d
Endosperm
Embryo
Anther
100%
Local Max
Linear
10700.3
9630.27
8560.24
7490.21
6420.18
5350.15
4280.12
3210.09
2140.06
1070.03
0
Masked (≥100% I

**(c)**

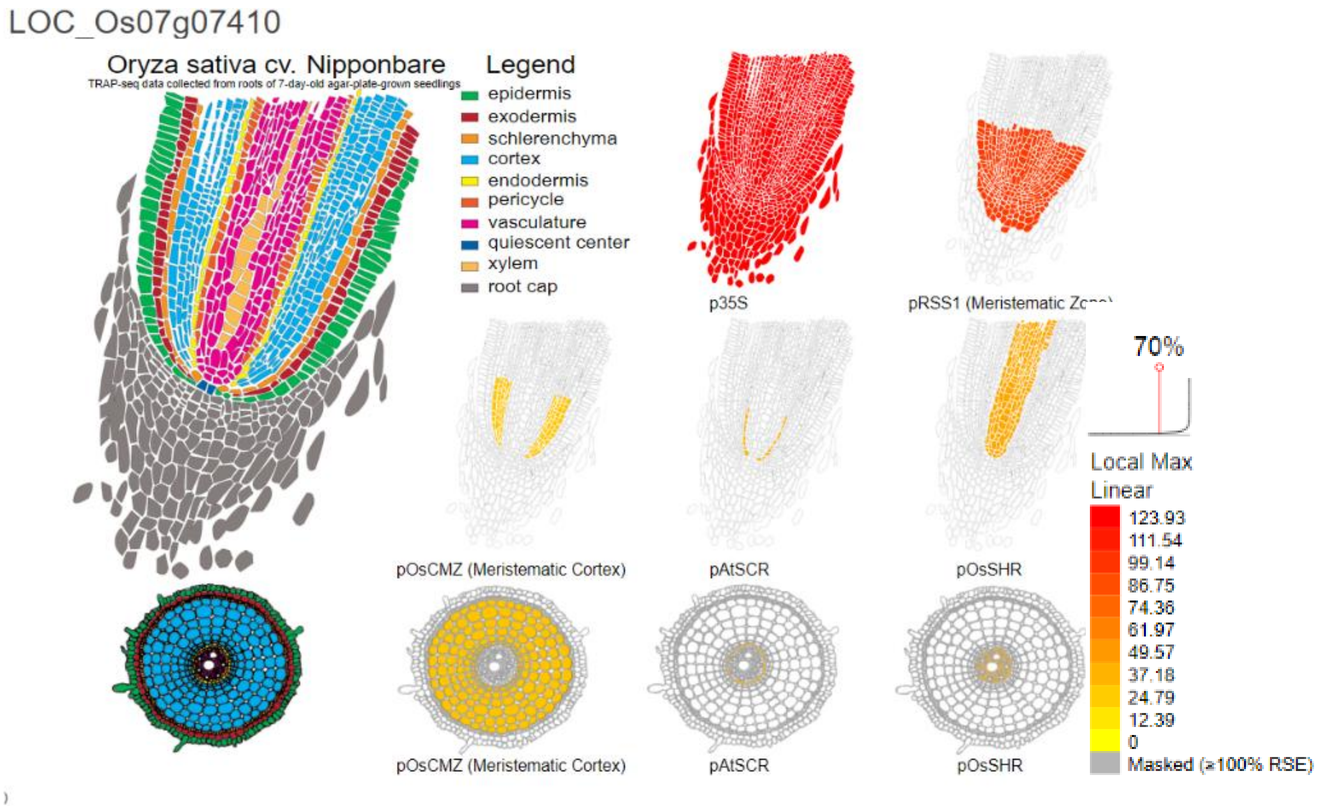


**Fig. S4** Spatiotemporal expression pattern of OsIDS3L. ePlant database showing the relative expression of OsIDS3L in different growth stage and tissues of rice plant. **(a)** Inflorescence, **(b)** Stigma and reproductive tissues and **(c)** Root tissues. The higher expression (red color) is observed dominant in roots, inflorescence and anthers. While lower expression was observed in other tissues (orange shades). The data curated from http://bar.utoronto.ca/eplant_rice/.

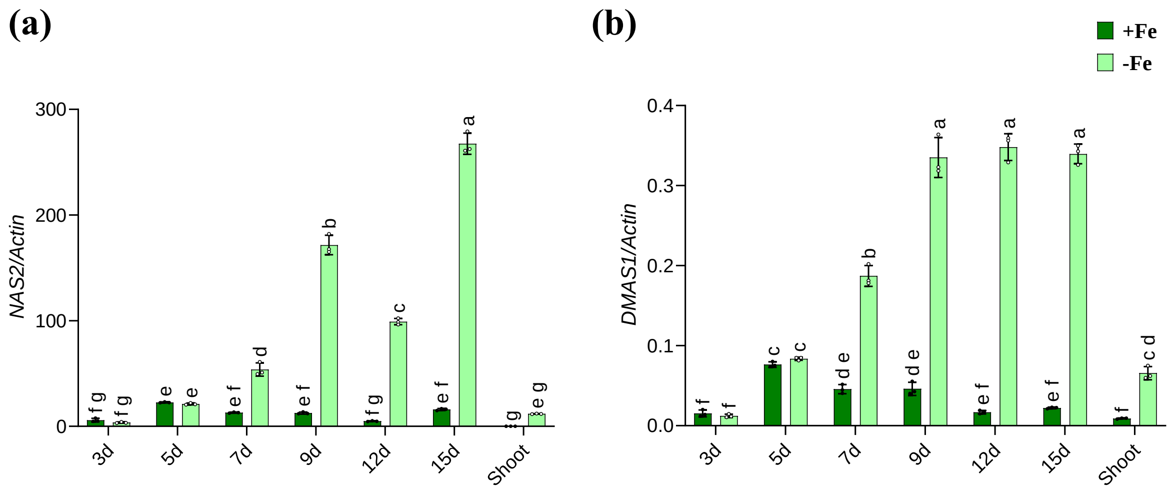


**Fig. S5** Fe deficiency response in rice at different stages. RT-qPCR expression analysis of **(a)** *NAS2* and **(b)** *DMAS1* in plant roots germinated and grown on ½ MS medium with or without 50µM NaFeEDTA (+Fe/-Fe) for the indicated number of days. Data represent mean ± SD from one biological replicate (n=3). Expression from 12-day-old shoot samples is presented as well. Similar expression was observed in other two biological repeats. Different lowercase letters indicate statistically significant differences as determined by a two-way ANOVA followed by Tukey's post hoc test.

**(a)**

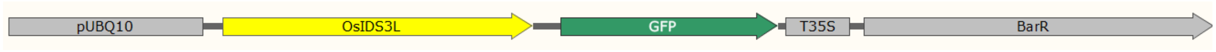


**(b)**

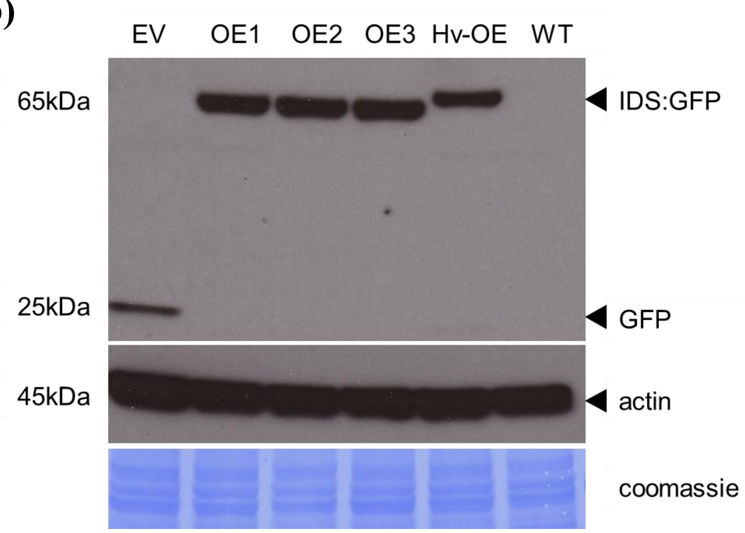


**Fig. S6** Generation of OsIDS3L overexpression lines **(a)** The vector construct used for OsIDS3L overexpression **(b)** Western blot of the transgenic lines against GFP antibody confirming induced transgene expression. Protein was extracted from 8-d old rice roots of respective lines grown and germinated on ½ MS media.

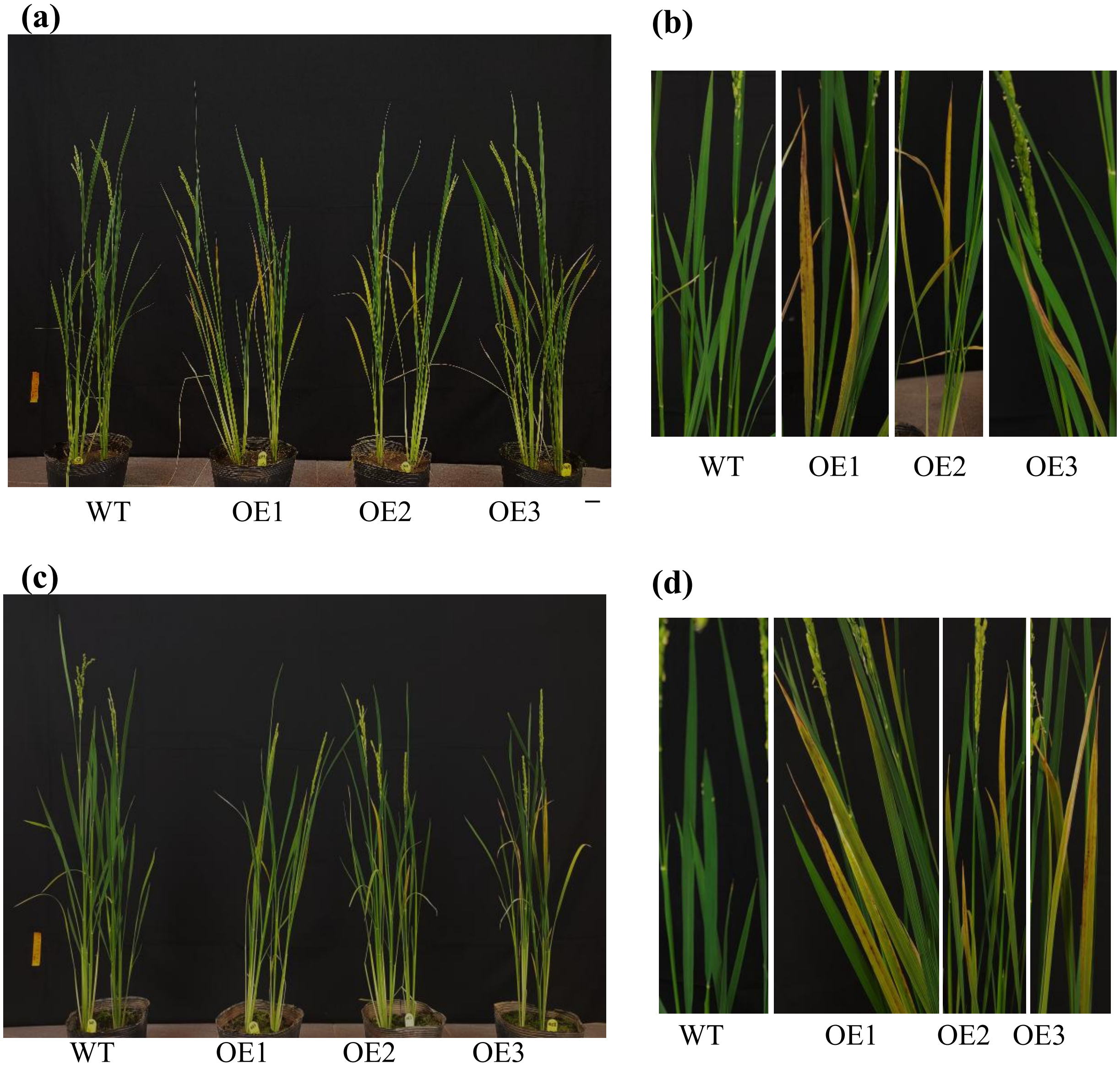


**Fig. S7** OsIDS3L overexpression lines shows bronzing symptoms when grown in soil. **(a-b)** water-logging condition **(c-d)** water-saving condition. The plant phenotype captured at early flowering stage. Bar =5 cm. **(b,d)** are enlarged leaves image depicting bronzing symptom.

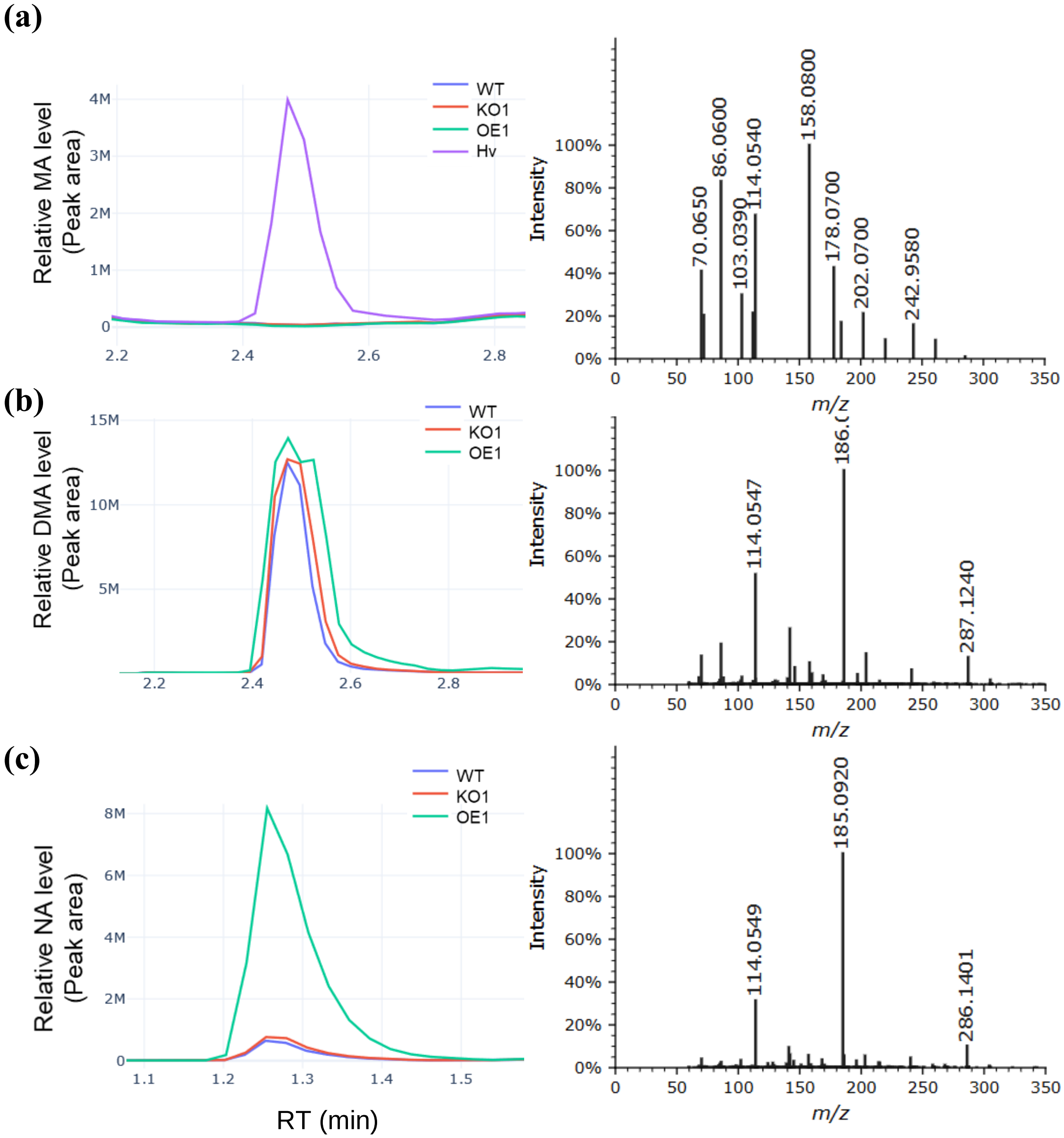


**Fig. S8** Extracted ion chromatogram (XIC) and the MS/MS fragmentation spectra of **(a)** MA **(b)** DMA and **(c)** NA. The images/data retrieved using GNPS2 dashboard. Data presented from metabolites extracted from plant roots germinated and grown under control conditions (1/2 MS with 50µM NaFeEDTA).

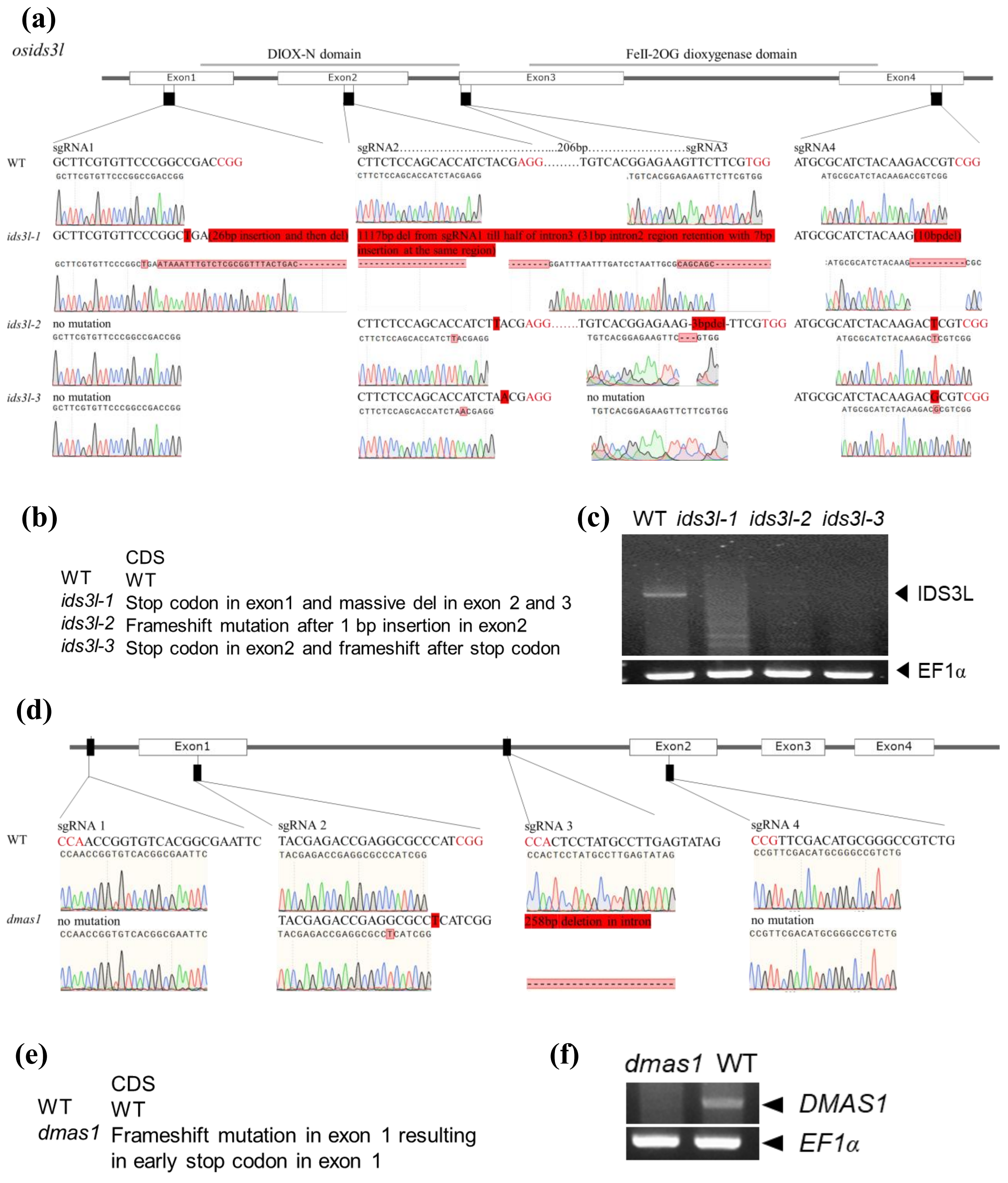


**Fig. S9** Generation of *ids3l and dmas1* knock-out lines. **(a)** Gene structure of OsIDS3L and the four sgRNA target sites used for CRISPR-Cas9 mutagenesis. Wild-type sequences are shown at the top; CRISPR-induced mutations in knockout lines are indicated in solid red boxes. PAM sequences (NGG) are highlighted in red. **(b)** CRISPR mutation effect on *OsIDS3L* CDS sequence. **(c)** Full-length RT-PCR of *OsIDS3L* in WT and three OsIDS3L-knock-out (*ids3l,* KO) lines. **(d)** Gene structure of OsDMAS1 and the four sgRNA target sites used for CRISPR-Cas9 mutagenesis. Wild-type sequences are shown at the top; CRISPR-induced mutations in knockout line are indicated in solid red boxes. PAM sequences (NGG) are highlighted in red. **(e)** CRISPR mutation effect on *OsDMAS1* CDS sequence. **(f)** Full-length RT-PCR of *DMAS1* in WT and DMAS1-knock-out (*dmas1*) lines.

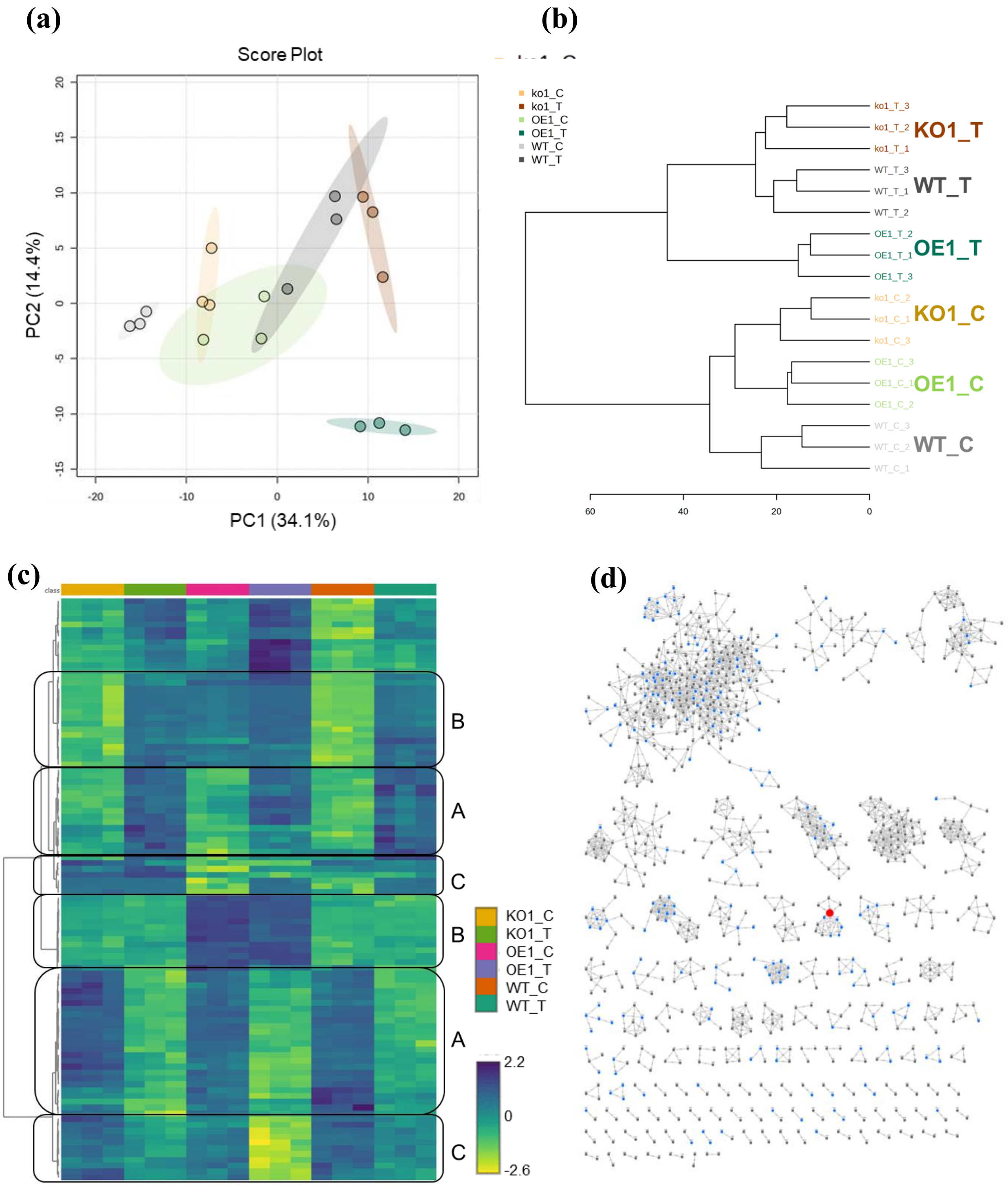


**Fig. S10** OsIDS3L transgenics shows significant metabolic perturbation. **(a)** Principal Component Analysis (PCA) of all metabolic features detected in untargeted metabolomics analysis. Grey, green and orange color are used to depict WT, OE and KO lines with darker shades showing Fe deficiency treated condition and lighter shades for control conditions. **(b)** Hierarchical clustering dendrogram showing sample segregation based on growth conditions and OsIDS3L enzyme expression. **(c)** Heatmap of 100 most variable metabolic features grouped into labeled blocks with distinct features. Group A shows metabolites responsive to Fe growth condition and not changed upon genotype. Group B and C shows metabolites affected by OsIDS3L gain- and loss of function (OE and KO). **(d)** GNPS-FBMN network showing the number of features and networks detected by aligning MS/MS fragmentation spectra. Highlighted red node represent a feature belonging to PS biosynthesis pathway components network.

**(a)**

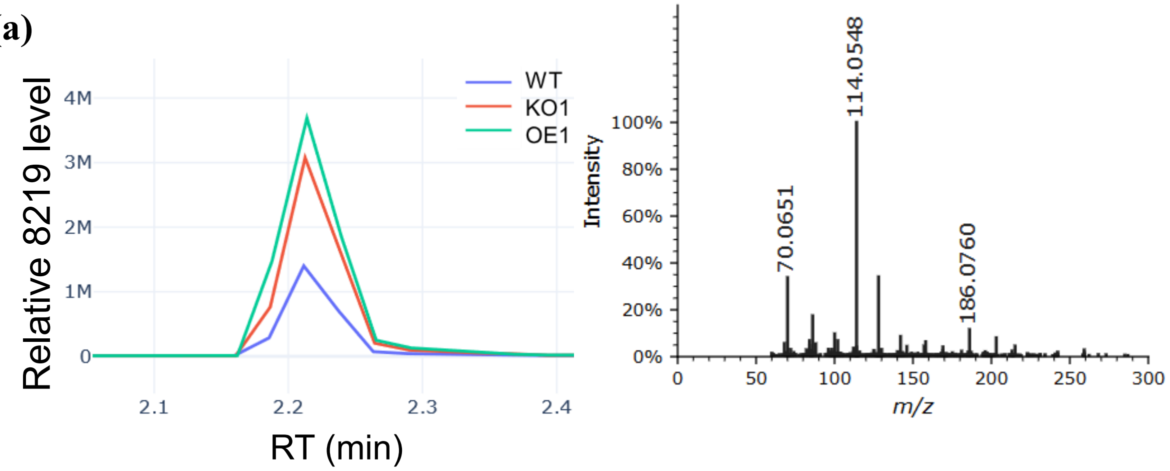


**(b)**

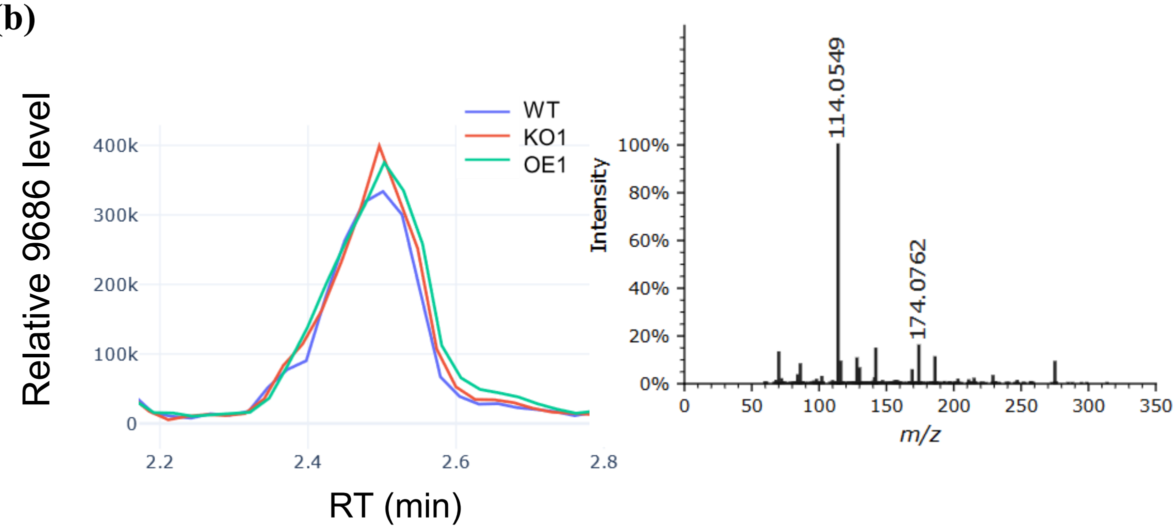


**Fig. S11** Extracted ion chromatogram (XIC) and the MS/MS fragmentation spectra of **(a)** 8219_275.1241_2.21 (depicted as, feature number_m/z_RT) and **(b)** 9686_259.1292_2.49. The images/data retrieved using GNPS2 dashboard. Data presented from metabolites extracted from plant roots germinated and grown under control conditions (1/2 MS with 50µM NaFeEDTA).

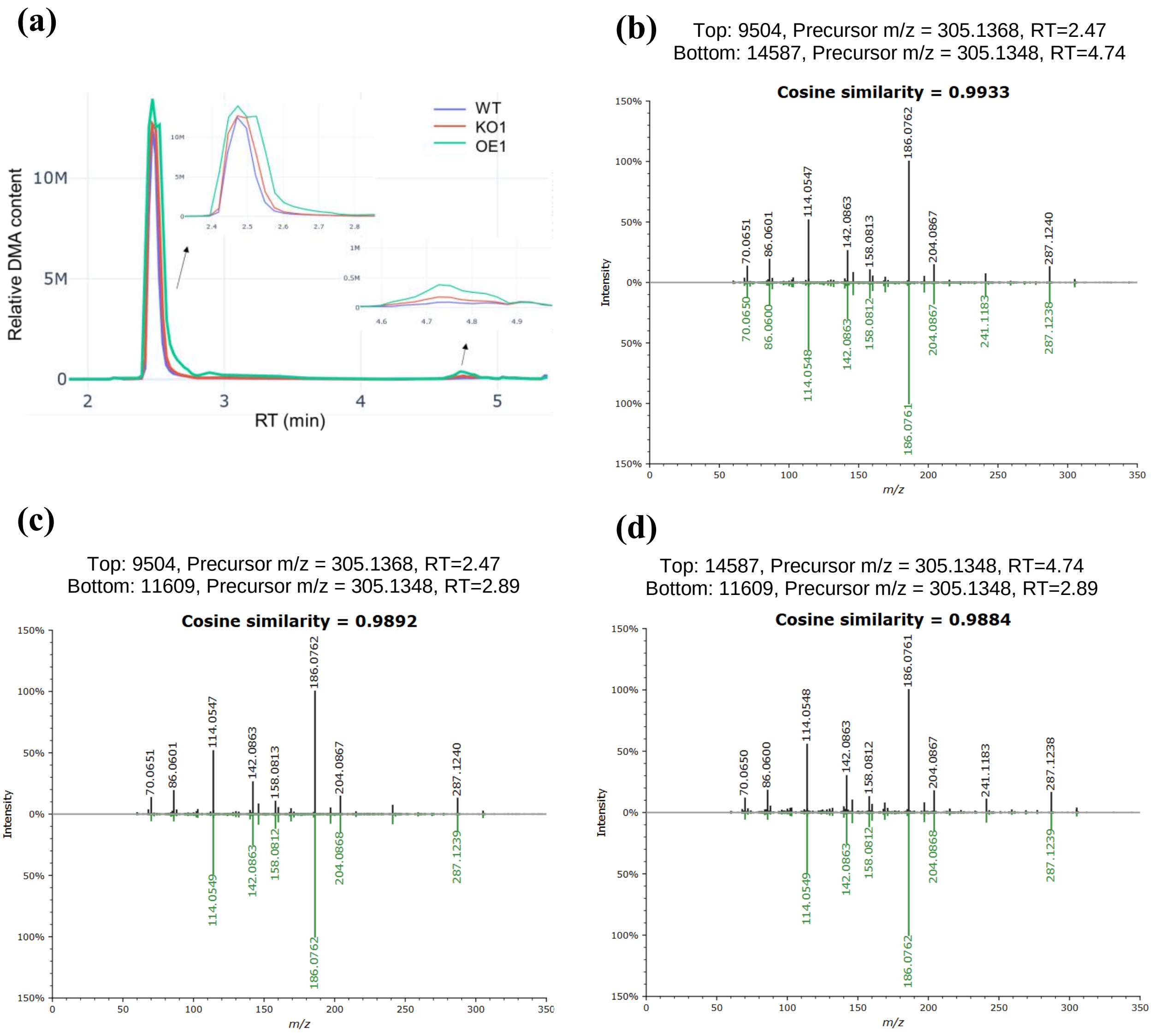


**Fig. S12** Feature and MS/MS fragmentation spectra highlight of three DMA nodes detected in GNPS-FBMN. **(a)** The DMA feature (305.13 m/z) at RT2.47, 2.89, and 4.74. Feature at RT 2.47 and 2.89 can be resolved to one feature based upon the broader elution time. The third feature at RT 4.74 hints stereoisomer. **(b-d)** Spectral similarity (MS2 spectra mirror plot) among three different node of DMA (nodes name: 9504, 14587, and 11609) showing a cosine score (MS/MS fragmentation spectra similarity score) higher than 0.95 indicating similar compound.

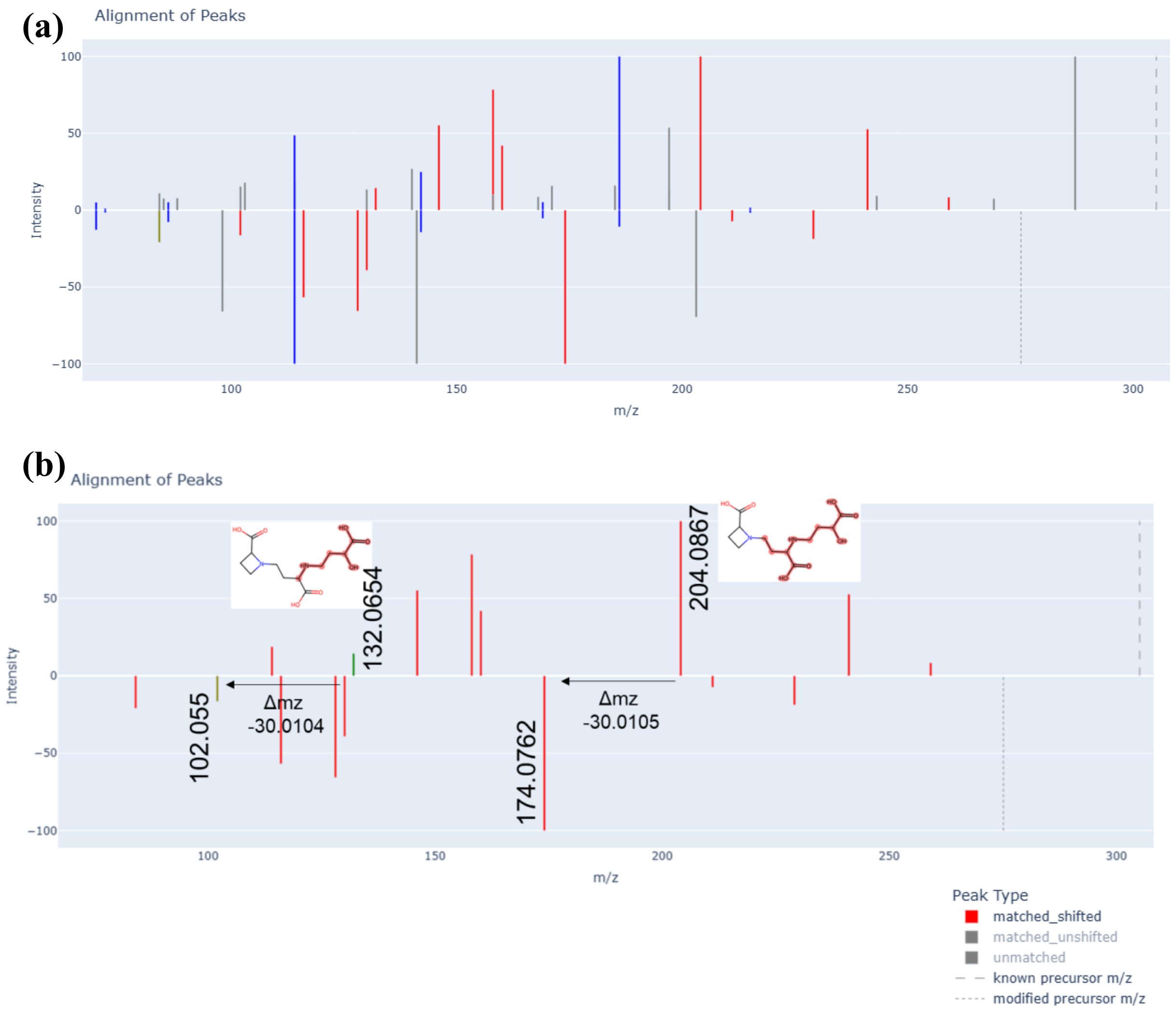


**Fig. S13** Structure prediction using Modifinder. The MS/MS fragmentation spectra was used to identify the structure of 8219_275.1241_2.21 (predicted product of OsIDS3L) using the known DMA structure. **(a)** The MS/MS spectra of DMA (top) and 8219 (bottom) represent precursor m/z in dashed line, matched unshifted peaks in blue and matched shifted peaks in red. **(b)** Match shifted peaks (red) were used to determine the possible site of modification and the structure. The MS/MS fragment shows the difference of 30 m/z in the side chain. The highlighted red structure of DMA shows the possible site of difference of $CH_2O$ group.

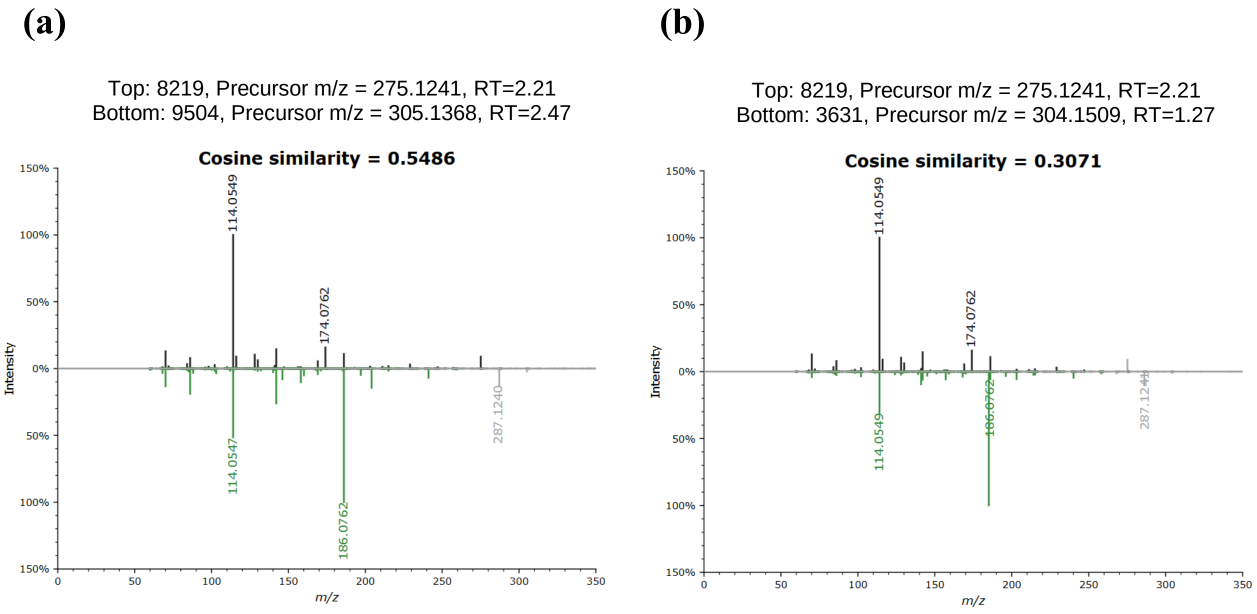


**Fig. S14** MS/MS fragmentation spectra similarity (cosine score) of metabolic feature 8219_275.1241_2.21 **(a)** with DMA, and **(b)** with NA.

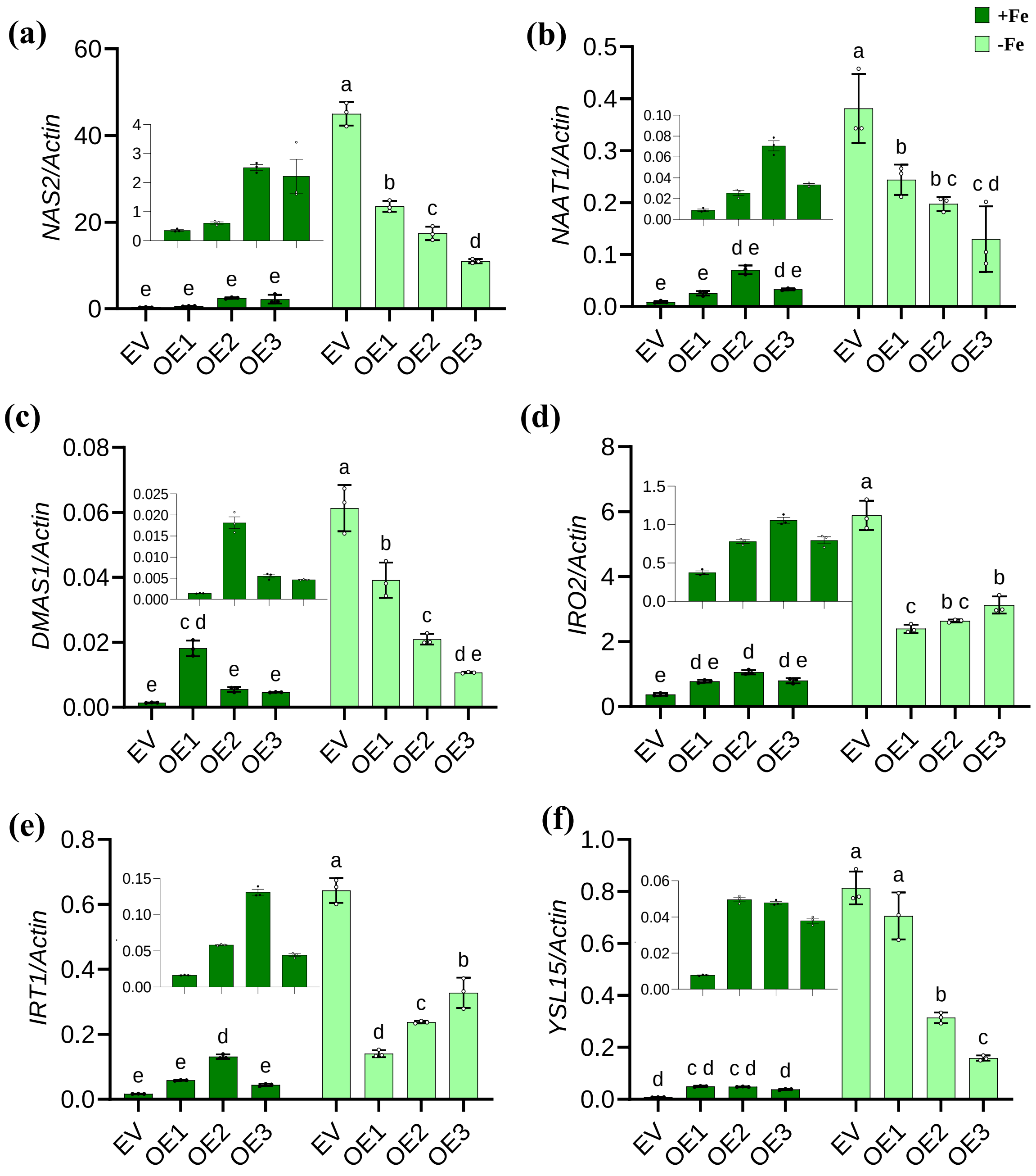


**Fig. S15** OsIDS3L alters baseline transcriptional response. RT-qPCR relative expression analysis of **(a)** *NAS2,* **(b)** *NAAT1,* **(c)** *DMAS1,* **(d)** *IRO2*, **(e)** *IRT1* and **(f)** *YSL15* in WT and OE roots. Plants were grown on ½ MS medium with or without 50µM NaFeEDTA (+Fe/-Fe) for 8 days. Data represent Mean ± SD, n=3. Different letters indicate significant differences determined by a two-way ANOVA followed by Tukey's post hoc test.
